# Antimicrobial effects, and selection for AMR by non-antibiotic drugs on bacterial communities

**DOI:** 10.1101/2024.04.23.590690

**Authors:** April Hayes, Lihong Zhang, Edward Feil, Barbara Kasprzyk-Hordern, Jason Snape, William H Gaze, Aimee K Murray

**Affiliations:** Faculty of Health and Life Sciences, University of Exeter; Department of Life Sciences, University of Bath; Department of Chemistry, University of Bath; Formerly AstraZeneca

**Keywords:** Antimicrobial Resistance, co-selection, non-antibiotic drugs, pharmaceutical pollution, metagenomics, environmental pollution

## Abstract

Antimicrobial resistance (AMR) is a major threat to human, veterinary, and agricultural health. AMR can be directly selected for by antibiotics, and indirectly co-selected for by biocides and metals. Some evidence suggests that non-antibiotic drugs (NADs) can co-select for AMR, but previous work focused on exposing single model bacterial species to predominately high concentrations of NADs. Here, we determined the antimicrobial effect and selective potential of three commonly used NADs against a complex bacterial community using a combination of culture based, metagenomic, and metratranscriptomic approaches. We found that three of five NADs tested on growth significantly reduced growth of a bacterial community, although only one (17-β-estradiol) selected for an AMR marker using qPCR. Whole metagenome sequencing indicated that there was no clear strong selection by NADs for antibiotic resistance genes, nor effects on community composition. However, some changes in relative abundance of metal resistance genes were observed after exposure to diclofenac, metformin, and 17-β-estradiol. Together, these results indicate that the NADs tested likely do not strongly select for AMR at both clinically and environmentally relevant concentrations.

## Introduction

Non-antibiotic drugs (NADs) are of increasing interest in terms of their effects on bacteria, including as potential antibiotic adjuvants to treat resistant pathogens (1). NADs can decrease the growth of both human gut strains (2), pathogens (3), and clinical and lab strains of bacteria (4–6). However, some NADs can also increase horizontal gene transfer (HGT) rates between bacterial species through transformation (7, 8), and conjugation (9–11), as well as selecting for antibiotic resistance through increasing mutation rates (12–15). Therefore, there are concerns that NADs may select for antimicrobial resistance (AMR). AMR is a significant threat to human health, with a predicted 1.27 million deaths already directly attributable to resistant bacterial infections in 2019 (16) and 10 million deaths estimated to occur annually by 2050 (17). Antibiotic resistance (occurring when bacteria become unable to be killed or inhibited by antibiotics) can be co-selected for by metals and biocides (18, 19). Co-selection refers to the simultaneous selection for resistance to two or more compounds (18). For example, exposing an aquatic bacterial community to benzalkonium chloride has been shown to increase resistance to ciprofloxacin and penicillin G (20). It is hypothesized that this could occur with NADs.

NAD concentrations are generally around the mg/L range in the human gut, (21) and around the ng/L – µg/L range in the aquatic environment (22, 23). Low concentrations of antibiotics and metals can select or co-select for antibiotic resistance genes (24–27), and these concentrations can be similar to those detected in the environment. Therefore, co-selection for AMR by NADs could potentially occur across a large geographic scale.

However, most current studies investigating NADs (to the authors’ knowledge) use single, model species of bacteria and mostly use high concentrations of NADs, e.g. g/mL or mg/mL concentrations, although there are some recent exceptions (e.g. (28)). It is unlikely that bacteria would be exposed to such high concentrations of NADs, with 20µM thought to be lower than the concentration found in the large or small intestine (2). Furthermore, single species experiments are unrealistic when considering bacteria exist in mixed species communities in nearly all known microbiomes. Mixed species models introduce social interactions (e.g. competition or co-operation) which has been shown to influence AMR (29, 30), and previous work has shown that the presence of a community can increase the selective concentration of gentamicin and kanamycin for a focal species (31). Therefore, in this study, a more realistic concentration range (high, gut relevant concentrations to low, environmentally relevant concentrations) was tested against a complex bacterial community.

The NADs tested in this study were diclofenac, ibuprofen, haloperidol, metformin, and 17-β-estradiol. These NADs can be detected in the aquatic environment (22, 32). Furthermore, all of these NADs have been shown to have antimicrobial effects, which may indicate potential to select for antibiotic resistance (33). Resistance to antibiotics commonly incurs a fitness cost, resulting in a reduction in the growth rate of the resistant strain relative to their sensitive counterpart ((33–37)). Therefore, it could be suggested that any reduction in growth with a compound (e.g. NAD) could indicate that resistance mechanisms may be required for survival/growth, and these could be co-opted for AMR. Previously, reduction in growth of a bacterial community has been shown to be a good proxy for selection for AMR markers, including *intI1* (33). Furthermore, modelling approaches have predicted that a reduction in growth rate is the most significant factor when attempting to estimate the minimal selective concentration (38).

The strongest evidence for antibiotic resistance co-selection by NADs exists for diclofenac, metformin and 17-β-estradiol. Diclofenac (a non-steroidal anti-inflammatory drug) can reduce bacterial growth (39–43), reduce biofilm formation (43, 44), and inhibit DNA synthesis (5, 45). Exposure to diclofenac can induce expression of genes that can increase AMR, increase resistance to antibiotics (14), and increase HGT (8). Furthermore, diclofenac is known to be a substrate of Gram-negative bacterial efflux pumps (39).

Metformin (a front-line treatment for Type II diabetes) can change the composition of the human gut microbiome (46, 47), and can reduce growth of bacteria (48–50). It has been demonstrated to increase mutation frequency and change gene expression, including AMR genes in *E. coli* (51). Metformin can also increase outer membrane permeability in *E. coli*, allowing for increased internal accumulation of tetracycline (50) and therefore potentially, other antibiotics.

The evidence regarding 17-β-estradiol (a natural hormone and used in hormone replacement therapy) is more conflicting. For example, it can both increase and decrease the growth of *E. coli* (52). *Prevotella intermedius* and *Prevotella melaninogenica* can use 17-β-estradiol as a vitamin K substitute, but at high concentrations, their growth was inhibited (53). It can increase soil biomass in some soils, but not in others, suggesting a soil microbiome/chemical specific response (54). 17-β-estradiol has also been shown to be a substrate of the multidrug efflux pump AcrAB-TolC (55), which confers resistance to multiple antibiotics (56)).

Two other NADs with some evidence of co-selection for antibiotic resistance were also tested. Ibuprofen (also a NSAID) can reduce the growth of bacteria (57), may alter the gut microbiome, can select for antibiotic resistance (58) and increase HGT (8). However, one study found no co-selective potential in *E. coli* after 30 day exposure to ibuprofen (28). It has also been shown to be a substrate of efflux pumps (39). Haloperidol (an antipsychotic medication) has only been demonstrated to reduce bacterial growth (2).

To determine if these NADs could co-select for AMR, a wastewater influent bacterial community was exposed to diclofenac, ibuprofen, haloperidol, metformin, and 17-β-estradiol separately. Effects on growth and selection for AMR were experimentally determined. For the former, a simple growth-based assay (33) was used to identify which NADs cause significant reductions in community growth. For the latter, phenotypic resistance was tested by plating NAD-evolved communities onto antibiotic amended agar plates. Further, DNA was extracted from the evolved communities and was used in qPCR assays (for *intI1, cintI1*, and *sul1*) and metagenomic sequencing. A general class I integrase gene (*intI1)* and a clinical class I integrase gene (*cintI1*) were used as gene targets, since *intI1* is often recommended as a gene target for AMR surveillance (59, 60), and because *intI1* can be associated with various antibiotic resistance gene cassettes (60) that can be selected for by antibiotics (27). Metatranscriptomic analyses were carried out on communities evolved for seven days with a high concentration of diclofenac, metformin, and 17-β-estradiol.

## Methods

### Inhibition of growth by NADs

Wastewater influent was collected from Falmouth, UK, WWTP, on the 2/2/2020, and frozen at 1:1 with 40% glycerol to create a final concentration of 20% glycerol and stored at −70 until use.

Diclofenac (Sigma Aldritch), ibuprofen (Sigma Aldritch), haloperidol (Sigma Aldritch), metformin (Enzo Life Sciences), and 17-β-estradiol (Sigma Aldritch) were dissolved in water, water, acetone, water, and ethanol respectively, and kept at −20°C before use, and used within nine days.

For all growth curve experiments, 5mL wastewater influent frozen stocks were thawed, and centrifuged at 14.8rpm for 2 minutes, before being washed with 0.85% NaCl (Sigma Aldritch) twice to remove nutrient and glycerol carry over from the frozen stock. The re-suspended pellet was used to inoculate Iso-Sensitest broth (Oxoid) at 10% volume (vol/vol). There were six replicates per treatment. There were no significant effects of solvent on growth of bacteria so all experimental runs only contained a no-NAD control. Concentration ranges of NAD started at 20µM, and then descended in a two-fold dilution until tested concentrations did not have any significant difference to the no-NAD control. All NADs were tested individually, i.e. no mixture effects were tested.

The 96-well plates containing 200µl growth cultures were incubated at 37oC in Varioskan Flash (Thermo Fisher) plate reader for 11 hours, with continual shaking at 180rpm. OD_600nm_ was taken every hour.

### Lowest Observed Effect (LOEC) Calculations

Were performed as previously described (33). In summary, the time point during exponential phase with the greatest dose response was determined using either a Spearman’s or Pearson’s correlation test. Then, optical density (OD) at all of the concentrations at that time point are tested to determine the concentrations that differ significantly from the control growth using a Dunn’s test (*dunn.test* v1.3.5). The lowest observed effect concentration (LOEC) was recorded as the lowest tested concentration at this time point that was significantly different to no-NAD control. The no observed effect concentration (NOEC) was recorded as the highest tested concentration that did not have any statistically significant effect on growth when compared to the no-NAD control (i.e., the test concentration directly below the LOEC). Ibuprofen and haloperidol did not significantly affect growth (see Results), and so were not tested further.

### Phenotypic antibiotic resistance

For phenotypic antibiotic resistance analysis, selection experiments only using 20µM of diclofenac (6400µg/L), metformin (3300µg/L), or 17-β-estradiol (5400µg/L) were performed. After seven days, 20µL of a dilution series of each of the five biological replicates for each treatment was plated onto Chromocult agar (Millipore) containing the EUCAST resistance endpoints for six antibiotics (Table 2) and spread across the plate using glass beads. A no-antibiotic control plate type was also used to determine the proportion of resistant colonies. Antibiotics were obtained: ampicillin (Melford), azithromycin (Merck), cefotaxime (Fisher), ciprofloxacin (Sigma Aldritch), gentamicin (Melford) and tetracycline (Formedium), and dissolved in water, ethanol, water, 0.5M hydrochloridic acid, water, and water respectively.

**Table 1.** Lowest Observed Effect Concentrations as determined through growth and *intI1* qPCR for diclofenac, metformin and 17-β-estradiol.

| Drug | Highest Concentration Tested ($\mu\text{g/L}$) | LOEC<sub>growth</sub> ($\mu\text{g/L}$) | LOEC<sub>qPCR</sub> ($\mu\text{g/L}$) |
| --- | --- | --- | --- |
| Diclofenac | 6400 | 50 | 0 |
| Metformin | 3300 | 26 | 0 |
| 17-β-estradiol | 5440 | 24.375 | 7 |

**Table 2:**
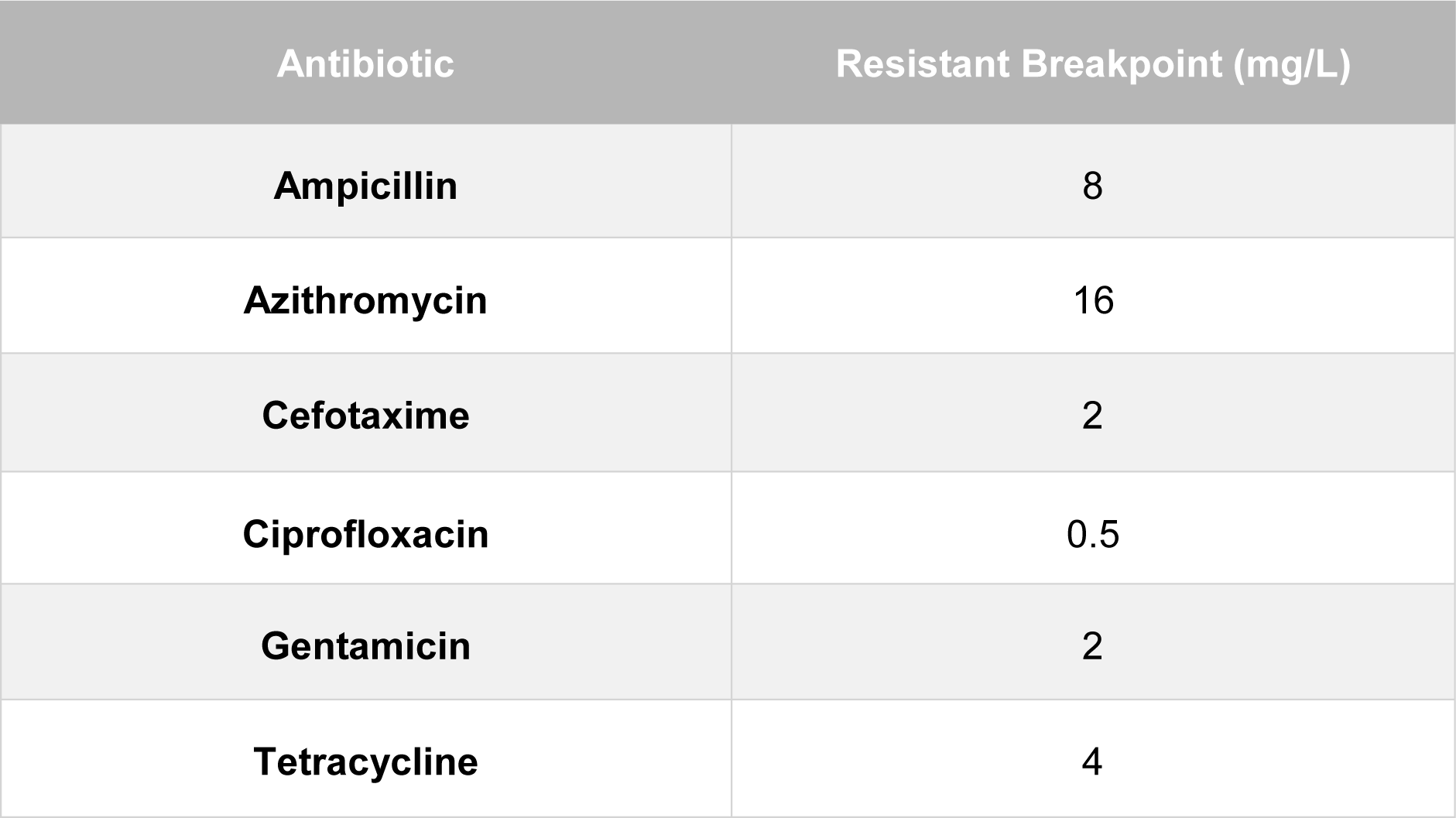
EUCAST breakpoints for *Enterobacterales* used for plating on Chromocult agar.

### Phenotypic resistance analysis

Colony forming units (CFU)/mL was calculated for all plate types. Resistant proportion was calculated by dividing resistant CFU/mL by total CFU/mL. Linear mixed effect models (*lme4*), with microcosm as a random effect, and antibiotic plate and NAD treatment as fixed effects were used to determine if NAD treatment changed CFU/mL on antibiotic amended plates. Models were run on all data (*E. coli* and presumptive coliform) and both of these separately. Fit of models was checked using *DHARMa.* Post-hoc tests were performed using *emmeans* to determine which treatments had significantly altered phenotypic resistance (CFU/mL).

### Longer-term selection of bacterial communities by NADs

Selection experiments were carried out using diclofenac, metformin, and 17-β-estradiol. Wastewater influent was thawed and washed with 0.85% NaCl, and spun at 3,500rpm for 10 minutes twice to remove contaminants and glycerol. The washed bacterial community was added to Iso-Sensitest broth (Oxoid) at 10% v/v. Concentrations of NAD, in a threefold dilution from 20µM to a concentration below the LOEC. All concentrations are listed. Diclofenac concentrations used: 6400µg/L, 2100µg/L, 710µg/L, 240µg/L, 79µg/L, 26µg/L, 8.8µg/L. Metformin concentrations used: 3300µg/L, 1100µg/L, 367µg/L, 122µg/L 40.7µg/L, 13.58µg/L, 4.53µg/L, 0µg/L. 17-β-estradiol concentrations used: 5440µg/L, 1813µg/L, 604µg/L, 201µg/L, 67µg/L, 22µg/L, 7µg/L, 2µg/L, 0µg/L. There were five replicates per concentration. Two 1mL aliquots were taken, spun down at 21,000 rpm for 2 minutes, re-suspended in 20% glycerol and frozen at −70°C until further use. The microcosms were incubated at 37oC with shaking at 180rpm, with 1% transfers daily to the same fresh medium. On day seven, four 0.5mL aliquots were taken and mixed with 40% glycerol (1:1), and frozen at −70°C until further use (qPCR, metagenomics and metatranscriptomics).

### QPCR

For qPCR, DNA was extracted from all day zero and day seven replicates using a Qiagen DNeasy UltraClean Microbial Kit, carried out to manufacturer’s instructions, with the initial centrifugation step increased to one minute. Extracted DNA was diluted 5x with 10mM Tris-HCl (Sigma Aldritch), and kept at 4oC until use in qPCR assays, or frozen at −20°C. If samples were thawed, they were kept at 4oC.

QPCR was carried out on all experiments using *intI1, cintI1* or *sul1* as a resistance endpoint, with normalisation to the 16S rRNA gene. DNA was diluted five times before use in qPCR, so that 5µL DNA added to the reaction contained 1µL undiluted DNA. The reaction mix contained 10µL PrecisionPlus Master Mix with ROX and SYBR, 1µL forward primer, 1µL reverse primer, 0.2µL bovine serum albumin (Fisher), and 2.8µL nuclease free water (Ambion). The reaction cycle was as follows: 120 second hold at 95oC, 40 rounds of cycling DNA amplification, with 10 seconds at 95oC and 60 seconds at 60oC for data collection. Only qPCR runs that had an R^2^>0.9 and an efficiency of between 90-110% were analysed. For *cintI1* assays, the annealing temperature was increased to 70oC. Primer sequences can be found in Table 3.

**Table 3:**
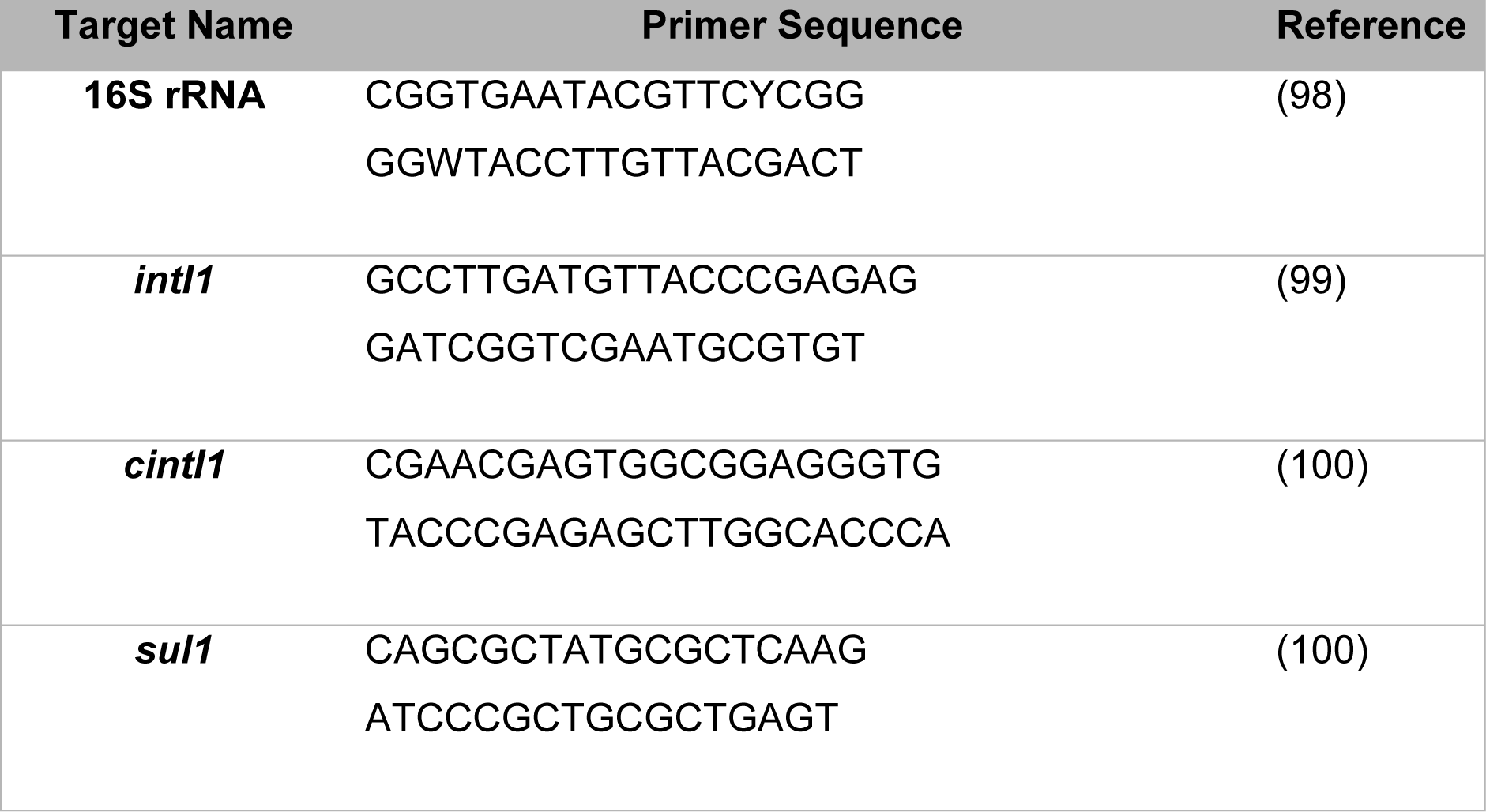
qPCR primer sequences.

Five concentrations of DNA standards were used for each gene to produce a standard curve. DNA standards (gBlocks) were purchased from IDT Technologies and sequences are presented in Table 4. Target gene molecular prevalence was determined by dividing the gene target quantity by the 16S rRNA quantity (i.e., *sul1* quantity/16S rRNA quantity).

**Table 4:**
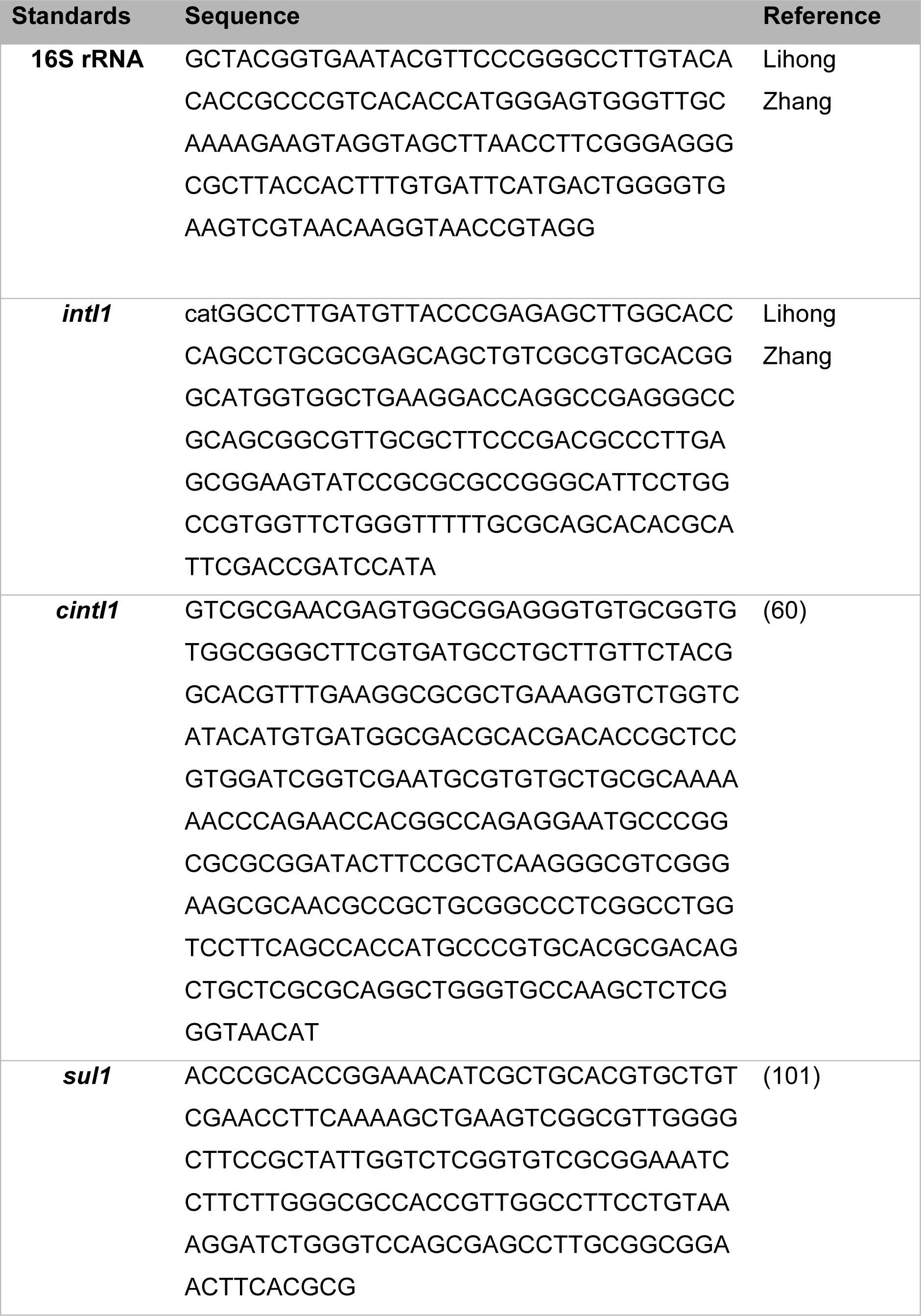
DNA standard sequences.

### QPCR analyses

Linear mixed effect models were run on the prevalence data using *lme4* v1.1.31 (61) in order to determine effect of treatment and time on the prevalence of *intI1, cintI1,* or *sul1* in the samples. The most parsimonious model was used, as determined using *DHARMa* v0.4.6 (62). Post-hoc comparisons were performed using *emmeans* v1.8.2 (63), which determined pairwise comparisons to assign significant selective concentrations.

### Metagenomic Sequencing

For metagenomic sequencing, DNA was extracted using the Qiagen DNeasy UltraClean Microbial Kit, and all steps carried out to manufacturer’s instructions, with the initial centrifugation step increased to one minute. DNA was extracted from three replicates for each day seven treatment, alongside three 40mL influent (inoculum) aliquots.

DNA was purified before being sent for sequencing using an Ampure XP protocol. This involved 50µL Ampure beads being added to the sample DNA, prior to incubation for five to ten minutes. Tubes were placed into a magnetic stand, which attracted the Ampure beads. When the liquid was clear, the supernatant was removed. The beads were delicately washed with fresh 80% ethanol, which was removed, then beads were resuspended with 10mM Tris-HCl (Sigma Aldritch) and incubated at 50oC for 10 minutes. The tubes were placed back into the magnetic stand for the incubation until the liquid was clear. The liquid (containing DNA) was removed and transferred into fresh tubes for use. DNA was sequenced by the Exeter Sequencing Centre using a NovaSeq SP to a depth of ∼ 5GB. The diclofenac and 17-β-estradiol experiments were sequenced using NEB Ultra ll FS library prep, and the metformin experiment used a PCR free library prep. This difference was due to changes in protocol at the ESS. Three inoculum samples were sequenced using each library prep to allow comparison within experiments.

### Community composition analyses

MetaPhlAn v2.0 (64) was used to determine community taxonomy. Forward and reverse reads were paired using FLASH2 (65) before being piped into MetaPhlAn. The composition of the community (and to determine species sorting effects) were analysed in two ways. Firstly, Kruskal-Wallis tests determined any change in any species detected in at least one sample, with adjustment for multiple testing using False Discovery Rate (*FDR*). Then, the beta diversity was determined using Bray Curtis dissimilarity distance, and visualised using a Non-metric MultiDimensional Scaling (NMDS) using *vegan* v2.5-7 (66) with a starting seed of 123. An Analysis of Similarity (ANOSIM) was used to determine differences in treatments using *vegan* using 9999 permutations to determine effect.

### Antibiotic gene analysis

ARGs-OAP v2.0 (67) was used to identify relative abundance of ARGs. The default parameters were used for ARGs-OAP (a percent identity of 95% or greater, matching 25bp or higher, with an evalue cut off of 1e-7). ARG hits were normalised to 16S rRNA reads that were generated using the first stage of the ARGs-OAP pipeline.

Antibiotic gene abundances were analysed by both antibiotic class, and total antibiotic resistance genes (i.e. all genes within class) using Kruskal-Wallis tests using all treatments, and p values were adjusted for multiple testing *FDR*. Diversity of ARGs was determined using Bray-Curtis dissimilarity, and visualised using NMDS plots. ANOSIM was used to determine differences in treatments.

For metformin *tolC* analyses, dose response curves were fitted using packaged *drc* (68). A four parameter log logistic curve was fitted with Box-Cox transformation, as this improved fit and heterogeneity of variance.

### Metal and biocide resistance gene analysis

Biocide and metal resistance gene (BMRG) changes were identified using BacMet (69). Hits to the BacMet database were filtered to 80% percent identity match of at least 25bp. Hits were subsetted into genes associated with plasmids, and genes associated with efflux using metadata extracted from the BacMet database. These genes were then tested using Kruskal-Wallis tests and *fdr* adjusted, and any genes with an adjusted p value<0.05 were then analysed using broad scale correlation tests, with adjusted p values. If genes were then significant, they were individually plotted. Diversity of BMRGs was determined using Bray-Curtis dissimilarity, and visualised using NMDS plots. ANOSIM was used to determine differences in treatments.

### Metatranscriptomics sequencing

For metatranscriptomics analysis, the selection experiments were repeated as for the phenotypic plating selection experiment, with five replicates each of a control (0µg/L NAD), 6400µg/L diclofenac, 3300µg/L metformin, and 5400µg/L 17-β-estradiol. On the last day, after the initial transfer, communities were incubated for six hours at 37oC with 180rpm shaking to allow communities to enter exponential phase. During this stage, 1mL samples were taken, pelleted, and the supernatant removed. Then, the samples (pellets) were freeze dried using 100% ethanol and dry ice. Once frozen, samples were stored at −70°C overnight until packaging for transport. Samples were transported on dry ice to Novogene in Cambridge. All RNA extraction, sequencing, quality control, and analysis were performed by Novogene. In brief, RNA was extracted from the samples, and quality checked. The samples were then sequenced using Illumina platforms to produce 1.9GB to 3.5GB raw data per sample. The data was quality checked and analysed by Novogene and presented in this paper.

### Metatranscriptomics analysis

All analyses of metatranscriptomics data was conducted by Novogene. Reads were assembled using Trinity for taxonomic, and gene expression analyses. Functional gene annotation was performed using Gene Ontology (GO) (70), Kyoto Encyclopedia of Genes and Genomes (KEGG) (71–73), evolutionary gene genealogy Non-supervised Orthologous Groups (eggNOG) (74), and Carbohydrate-Active Enzymes (CAZy) (75) databases. Differential gene expression analysis was performed by mapping sample reads to assembled transcriptome and converting read counts to fragments per kilobase of transcript sequenced per million base pairs sequenced. Read counts were normalised, and multiple hypothesis test corrections used to produce adjusted p values. Reads were assembled using Trinity (76) by Novogene, and analysed to determine taxonomic changes, and functional gene analyses by Novogene. Species annotation was carried out using DIAMOND (77). Gene expression analysis mapped reads to the KEGG database. Differentially expressed genes (those with a >log2 fold change) between NAD treated communities and control communities was also determined by Novogene.

The reads for each sample were also piped through the ARGs-OAP and BacMet pipelines by the authors to determine changes in expression of these genes. The data from these were analysed as for the metagenome analyses.

### Statistical Analysis

All statistical analyses were carried out in R v4.1.10 (78). All plots were made using *ggplot* v3.3.5 (79). *Metbrewer* v0.2.0 (80) was also used to produce the plots.

## Results

### Three non-antibiotic drugs decrease growth of a bacterial community over 12 hours

Firstly, we investigated whether the NADs could reduce bacterial growth at a range of concentration, and aimed to identify the lowest observed effect concentration (LOEC) of each. We found that diclofenac, metformin, and 17-β-estradiol significantly decreased the growth of a bacterial community in exponential phase over 12 hours. The LOECs are shown in Table 1, and growth curves in Figure 1. Ibuprofen and haloperidol did not significantly reduce the growth of the community over 12 hours (p>0.05). Since only diclofenac, metformin, and 17-β-estradiol reduced the growth of the community, only these three NADs were tested further for their potential co-selective effects.

**Figure 1.**
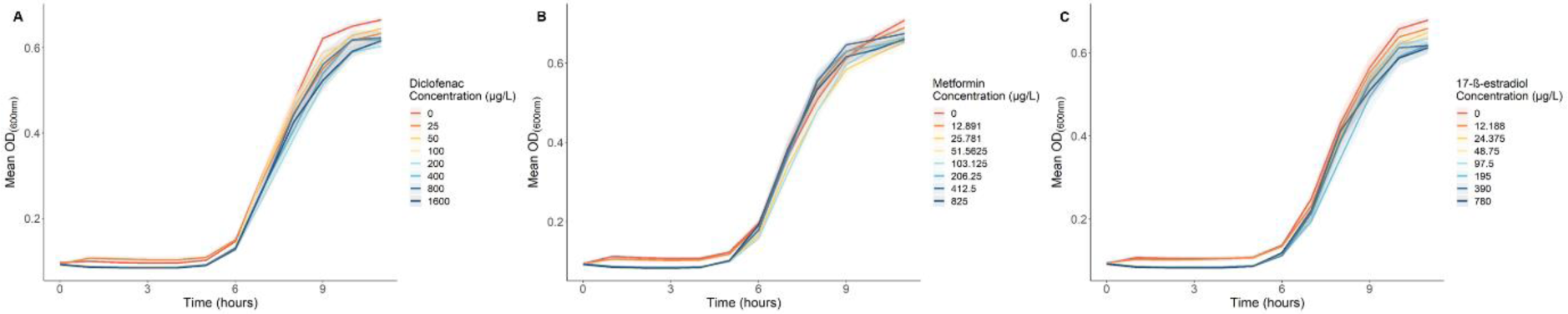
Growth over 11 hours of a mixed bacterial community with diclofenac (A), metformin (B), and 17-β-estradiol (C). Means of six replicates is shown, with pale ribbon standard error.

### Diclofenac, metformin, and 17-β-estradiol did not select for phenotypic resistance

To test for phenotypic expression of ARGs, cultures were plated onto coliform selective antibiotic amended plates after seven days evolution. When both *E. coli* and presumptive coliform CFU/mL were combined, NAD treatment did not significantly affect CFU/mL counts (NAD treatment main effect, (X^2^=6.85, d.f.=3, p=0.077). There was a similar response seen in the presumptive *E. coli* only data, with NAD treatment not significantly affecting the data (X^2^=1.95, d.f.=3, p=0.58). When only presumptive coliform counts were analysed (Figure 2), the interaction between NAD treatment and antibiotic plate, is significant (p=0.029). However, when tested, only the non-interacting terms of NAD treatment (NAD main effect: X^2^=10.74, d.f.=3, p=0.013), and type of antibiotic plate (antibiotic main effect: X^2^=84.32, d.f.=5, p=<0.0001) significantly affected the proportion of resistant CFU/mL. After *fdr* adjustment, pairwise comparisons show that 17-β-estradiol decreased resistance (i.e., resulted in collateral sensitivity) to ampicillin (p=0.0306) and gentamicin (p=0.017).

**Figure 2.**
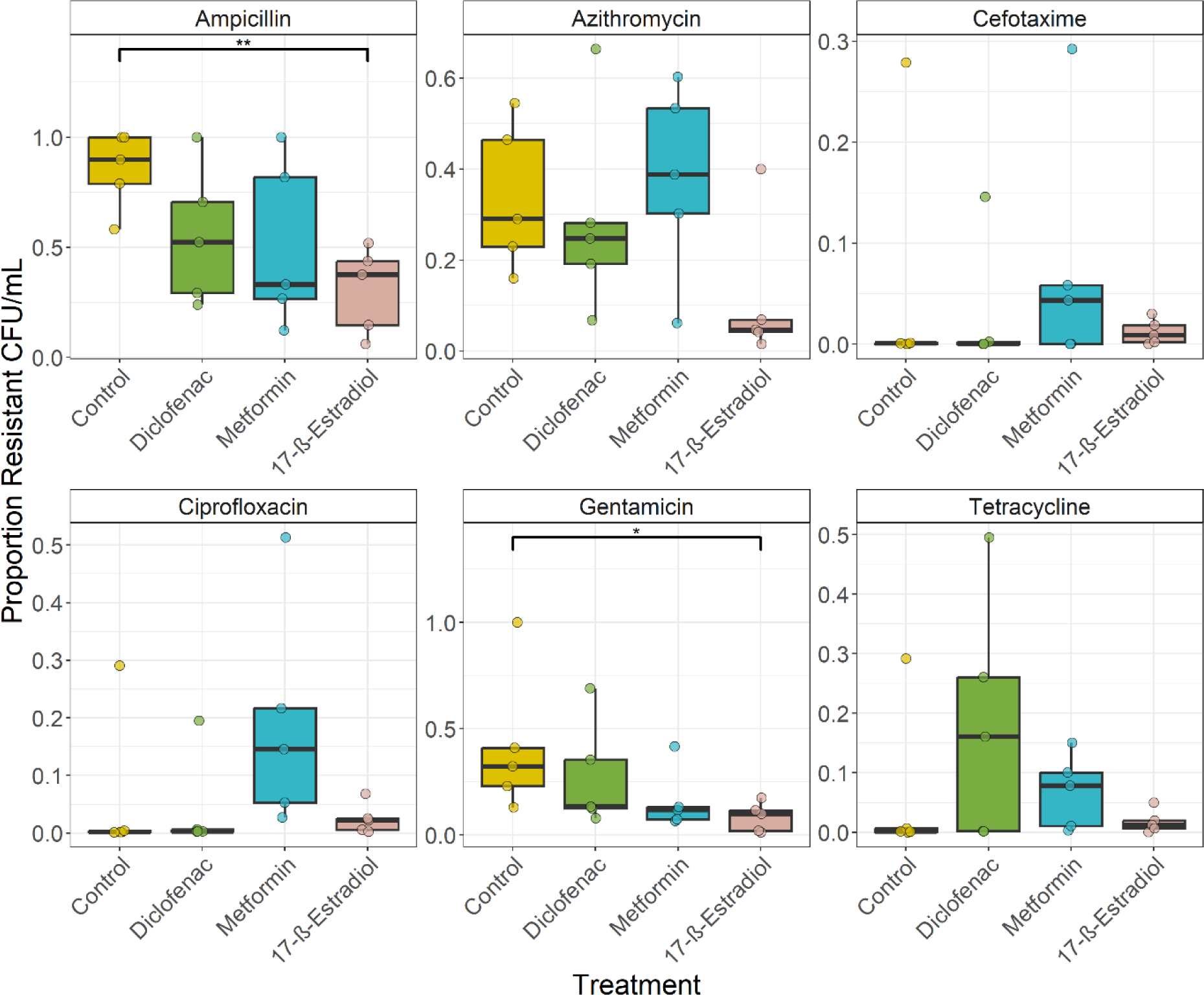
Resistant proportion of colony forming units (CFU) per mL growth on different antibiotic plates, with treatment with diclofenac, metformin, and 17-β-estradiol, presumptive coliform growth only.

### 17-β-estradiol selects for intI1, diclofenac and metformin do not

To identify common ARGs associated with antibiotic resistance, qPCR assays for *intI1, cintI1* and *sul1* were performed on the seven day samples. Firstly, *intI1* prevalence (*intI1* copy number normalised to 16S rRNA copy number) after exposure to diclofenac, metformin, and 17-β-estradiol was determined. *IntI1* was used as a general marker for AMR since it was unclear which AMR genes (if any) would be selected for after exposure to NADs. We found that diclofenac and metformin did not select for *intI1* after exposure for seven days (Supplementary Figures 1 and 2). However, 17-β-estradiol did select for *intI1* after exposure for seven days (Figure 3). Prevalence of *intI1* increased with time (time main effect: X^2^=103.31, d.f.=1, p<2.2e-16), and 17-β-estradiol concentration (X^2^= 29.76, d.f.=8, p=0.00023), but the interaction between the two was not significant. Pairwise comparisons showed that 7µg/L (p=0.0004), 22µg/L (p=0.0026), 67µg/ (p=0.0017), 201µg/L (p=0.0006), 604µg/L (p=0.0027), 1800µg/L (p=0.0050), 5400µg/L (p=0.0008) all significantly increased *intI1* prevalence compared to the control *intI1* prevalence at day seven.

**Figure 3:**
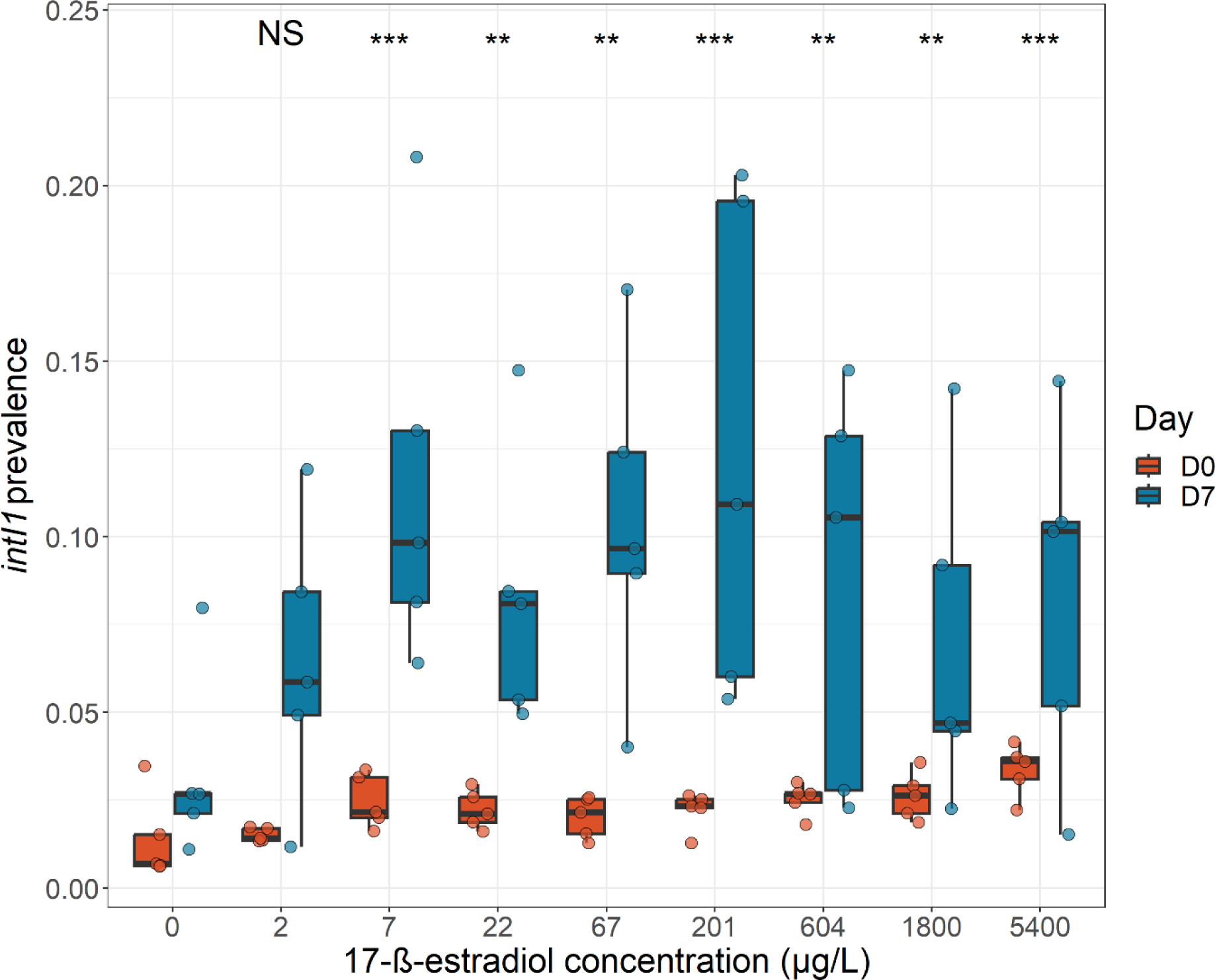
*IntI1 p*revalence for selection experiment with 17-βestradiol. 5 biological reps per treatment. Significance indicates significant difference in day seven prevalence compared to day seven prevalence at 0µg/L when analysed after pairwise comparisons *fdr* adjustment. Significance values as follows: NS - non-significant, *-P<0.05, ** - p<0.01, ***-p<0.001.

*CintI1* and *sul1* were not selected for by any of the three NADs, but did increase in prevalence with time (Supplementary Figures 1-3). Since the qPCR assays indicated that only 17-β-estradiol selected for a commonly used AMR marker, but all three NADs significantly reduced the growth of the community, full PCR-free metagenome sequencing was used to try to identify if changes to other AMR genes occurred after exposure to the NADs. Sequenced metagenomes were analysed to identify changes to community composition, antibiotic resistance genes (ARGs) and metal and biocide resistance genes (BMRGs).

### There was no ecological species sorting effects with NAD treatment

Overall, there was no ecological species sorting effects seen after exposure to NADs. Firstly, there was no significant selection for any detected bacterial species after treatment with either diclofenac, metformin, or 17-β-estradiol (Kruskal-Wallis, p>0.05, *fdr* adjustment). Secondly, there was also no change in the beta diversity of evolved populations treated with either diclofenac, metformin, or 17-β-estradiol (diclofenac, ANOSM, statistic R=0.0069, p=0.46, metformin ANOSIM, statistic R=-0.023, p=0.56, 17-β-estradiol ANOSIM, statistic R=0.12, p=0.94).

### There was non-specific selection for ARGs in evolved communities with treatment by diclofenac, metformin, or 17-β-estradiol

Firstly, we found that diclofenac and 17-β-estradiol did not significantly select for any ARGs detected (Kruskal-Wallis, p>0.05, *fdr* adjustment), and neither significantly altered the diversity of the ARGs within the evolved communities (diclofenac ANOSIM statistic R=0.012, p=0.054, 17-β-estradiol ANOSIM, statistic R=0.12, p=0.094, Supplementary Figures 7 and 9).

Although metformin did not specifically change the abundance of any antibiotic gene class (Supplementary Figure 8), metformin treatment of 13.6µg/L to 3300µg/L decreased diversity of ARGs compared to 0µg/L and 4.5µg/L evolved communities (Bray Curtis distance, ANOSIM, statistic R=0.374, p=0.0016).

Additionally, metformin treatment appeared to select against the efflux pump gene *TolC* (Kruskal-Wallis, p=0.026) (Figure 4) in a dose-response manner. The associated efflux genes *acrA* and *acrB* also showed decreased abundance, but this was not significant (Kruskal-Wallis, p=0.08, and p=0.051 respectively).

**Figure 4.**
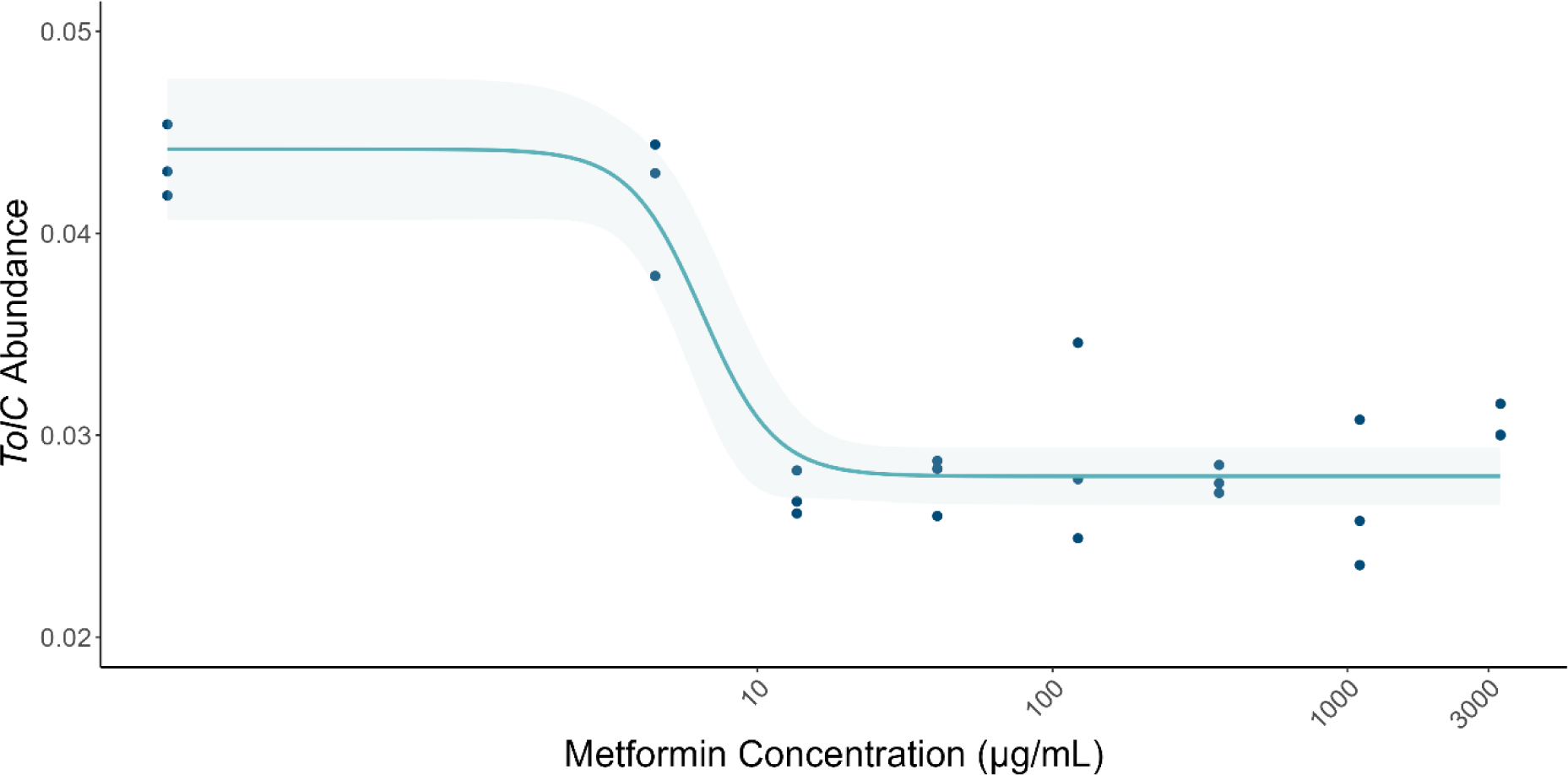
Relative abundance of *tolC* as a function of metformin concentration. Four parameter log logistic curve plotted (line) with 95% confidence intervals (pale ribbon) against all data points.

### All three NADs did select for some BMRGs

Diclofenac, metformin, and 17-β-estradiol individually, did not significantly change the relative abundance of any BMRG when all genes were tested together (Kruskal-Wallis, p>0.05, *fdr* adjustment). However, diclofenac caused a significant difference in the beta diversity of BMRGs between the evolved populations (ANOSIM statistic R=0.26, p=0.012). These differences may have been non-specific, since the communities did not become more dissimilar the more they varied with concentration (Mantel statistic R=-0.012, p=0.51). Metformin showed a non-significant change in the beta diversity of BMRGs (ANOSIM statistic R=0.17, p=0.054). There was also no significant change in the beta diversity of BMRGs in communities evolved with 17-β-estradiol (ANOSIM statistic R=0.077, p=0.19).

We tested the BMRGs that were specifically plasmid-borne, since mobile genes are more likely to be horizontally transferred between bacterial cells. Further, plasmid-borne genes are more likely to be co-located with multiple different genes that could confer resistance. Diclofenac treatment significantly altered relative abundance of 57 plasmid associated genes (including arsenic, mercury, nickel and tellurium resistance genes, Supplementary Table 1. However, these were not altered in a dose-dependent manner (Kruskal-Wallis test, p>0.05, *fdr* adjustment). Metformin treatment significantly altered relative abundance of 133 plasmid associated genes (Supplementary Table 2), again, in a non-dose-dependent manner (Kruskal-Wallis, p<0.05, *fdr* adjustment).

When populations were treated with 17-β-estradiol, there were 142 plasmid associated genes that were significantly different in at least one treatment (Kruskal-Wallis, p<0.05, *fdr* adjustment, Supplementary Table 3), but 138 were not linear dose-dependent. However, relative abundance of four genes significantly correlated with 17-β-estradiol concentration (Figure 5). We found that 17-β-estradiol positively selected for two copies of *arsB* and for *arsR*, but selected against *ncrA*.

**Figure 5.**
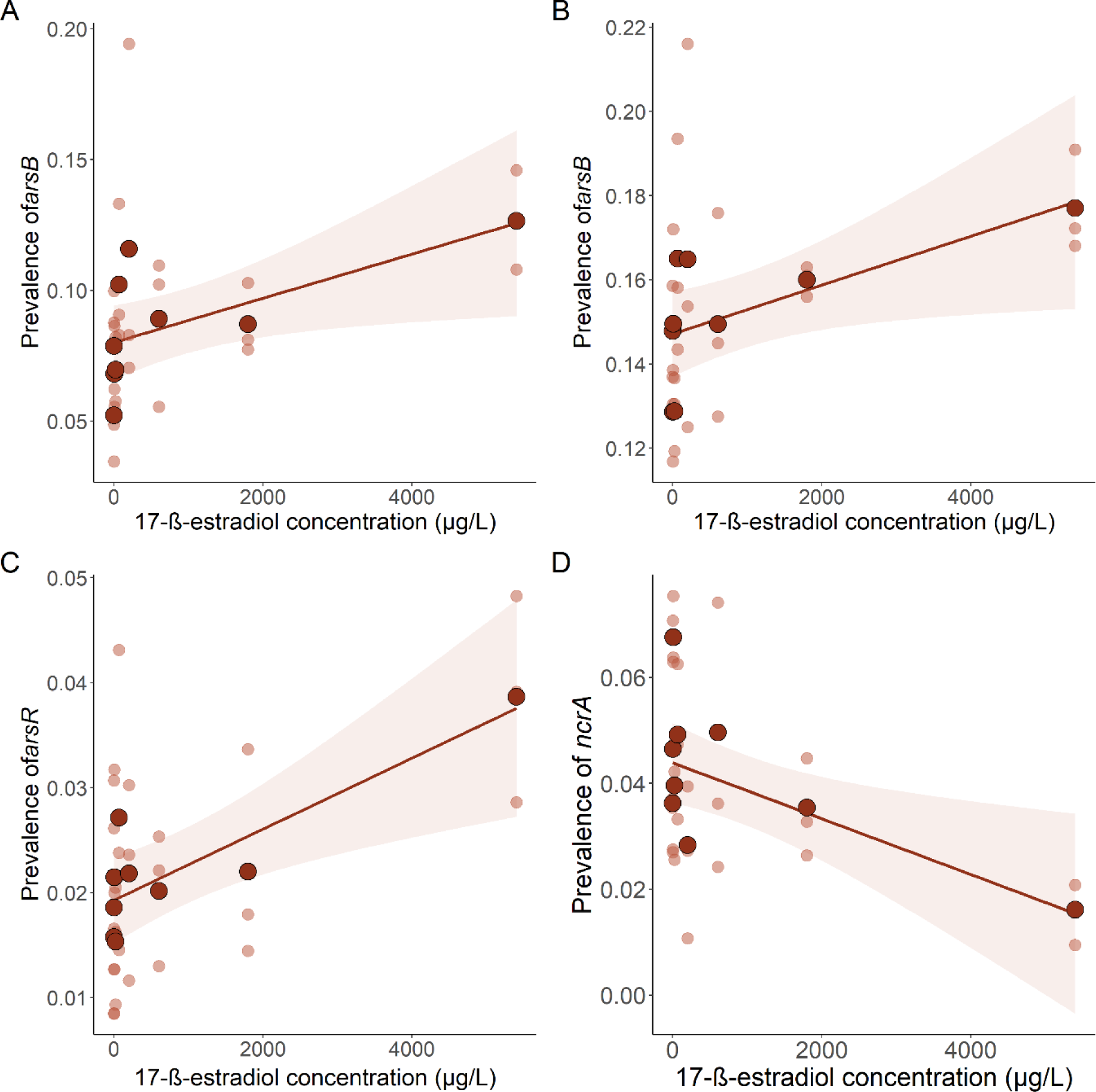
Genes that are significantly up or down correlated with 17-β-estradiol concentration. A – *arsB*, B – *arsB,* C – *arsR*, D – *ncrA*.

### NADs did not change ARG expression levels when tested at high concentrations

Since we did not find any specific selection for heritable ARGs or BMRGs that could explain the reduction in growth that we found, we decided to investigate the gene expression of communities that had been grown in a high concentration of NAD for seven days.

We did not find any ARGs (according to transcripts run through ARGs-OAP v2.0 (67) that were differentially expressed when communities were grown at high concentrations of diclofenac, metformin, or 17-β-estradiol when compared to the control. Additionally, there was no change in the functional gene pathway expression (e.g., KEGG) that these communities were expressing (Kruskal-Wallis, p<0.05, *fdr* adjustment). However, there were several thousand unidentified genes that were only expressed in either the diclofenac (n=8,967), metformin, (n=11,063), or 17-β-estradiol (n=9,494) treatments and not the control, with a further 1,081 unidentified genes expressed in all of the NAD treatments but not the control populations.

#### Antibiotic Resistance Genes

The expression of *acrA*, *acrB*, and *TolC* genes were specifically investigated since it has been previously suggested that diclofenac, metformin, and 17-β-estradiol are acrAB-TolC efflux pump substrates (39, 51, 55). However, there was no significant change in the expression of these genes resulting from NAD treatment (Kruskal-Wallis, *acrA* p=0.65, *acrB* p=0.61, *TolC* p=0.83). Additionally, there were no significant changes in expression at the ARG class level, following NAD treatment (Kruskal-Wallis, p<0.05, *fdr* adjustment). There was also no significant difference in beta diversity of the expressed genes between NAD treatments for both ARG classes (ANOSIM statistic R=0.04, p=0.26), and all ARGs (ANOSIM statistic R=0.008, p=0.41).

### NADs may change BMRG expression levels

NAD treatment did not significantly affect the expression of BMRGs involved with efflux (Kruskal-Wallis, p>0.05, *fdr* adjustment). However, there was a significant change in the expression of plasmid associated genes. Five plasmid associated BMRGs were significantly different across the NAD treatments (Figure 6, Kruskal-Wallis, p<0.05, *fdr* adjustment). We found that diclofenac reduced the expression of *arsR* (Dunn’s test, p=0.024), metformin reduced the expression of *arsB* (Dunn’s test, p=0.014), but was shown to increase the expression of *merP* (Dunn’s test, p=0.044). Finally, 17-β-estradiol reduced the expression of *arsR* (Dunn’s test, p=0.031) and *merA* (Dunn’s test, p=0.013) at this high concentration. However, there was no overall significant change in the beta diversity of the expressed BMRGs between the NAD treatments.

**Figure 6.**
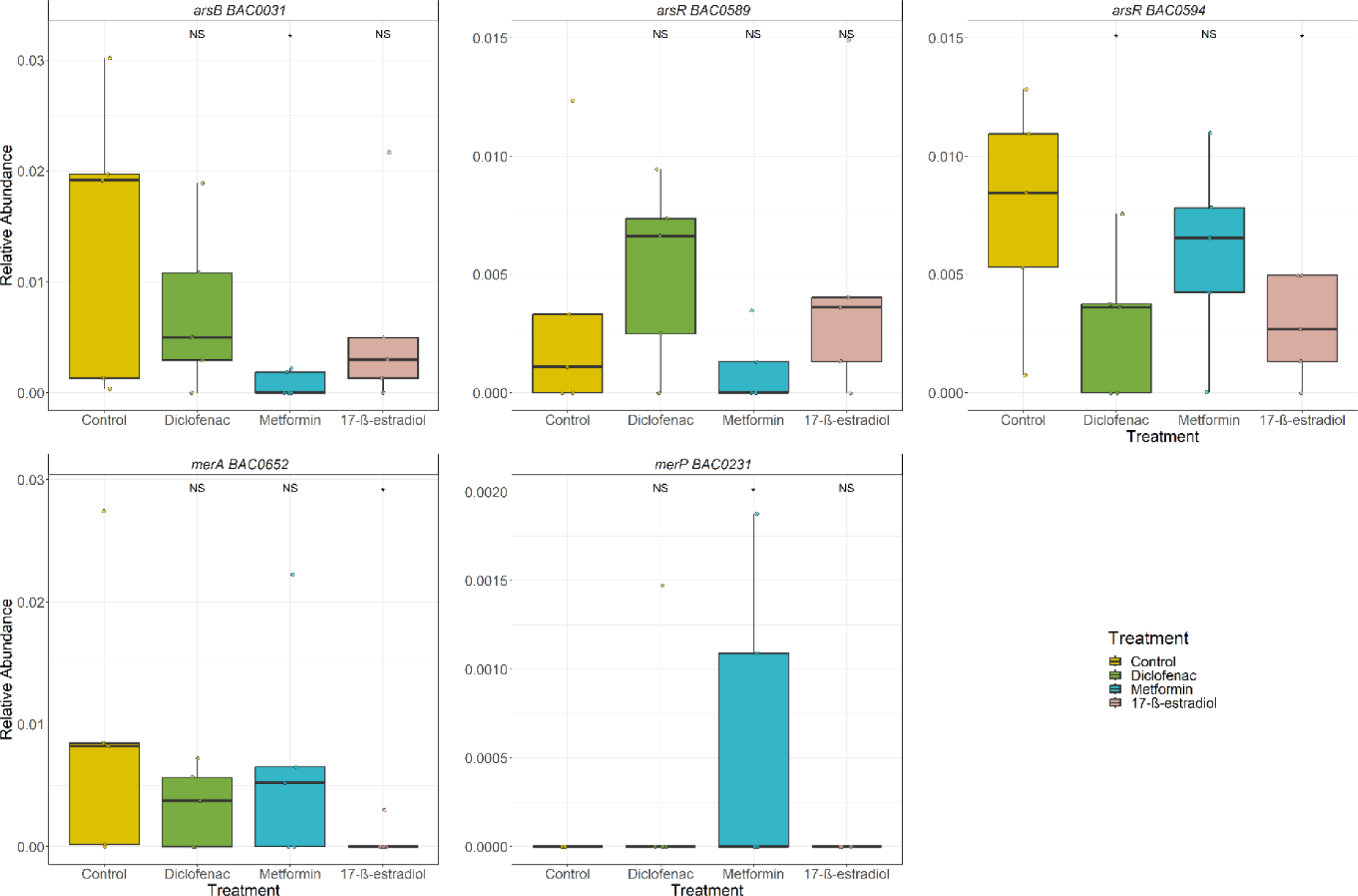
Expressed BMRGs in communities evolved with diclofenac, metformin, or 17-β-estradiol, or a no-NAD control. P<0.05 = *, p<0.01 = ** (Dunn’s test).

## Discussion

The effects of five NADs were tested on a wastewater influent bacterial community. Only three tested NADs significantly reduced the growth of the community, and were tested further. Ibuprofen and haloperidol did not significantly reduce growth of the wastewater influent bacterial community here, although they have previously been indicated to have growth inhibitory effects (2, 57). These differences may be due to different experimental designs. For example, in this study we used a mixed bacterial community derived from wastewater, whereas much previous work has focused on single species models (e.g. (2, 3)).

### Short term effects on growth and community functioning

Diclofenac, metformin, and 17-β-estradiol affected growth as concentrations similar to those found in the aquatic environment, both globally (22) and in England and Wales (81). This suggests that in the environment, diclofenac, metformin, and 17-β-estradiol may alter the functioning of the bacterial populations present there, indicating that increased pollution via wastewater into the freshwater environment may affect ecosystem functioning. WWTPs rely on microbial functioning during activated sludge treatment, and the ability of the community to degrade organic material varies with different stresses, and this may also be the case for these pharmaceutical pollutants (82–86). Additionally, the various gene transcripts only expressed in NAD treatments could also be associated with changes to community functioning, and further work should aim to identify changes to housekeeping functions under NAD stress.

### Longer term effects on AMR

The longer-term effects of diclofenac, metformin and 17-β-estradiol were tested on the same type of wastewater influent bacterial community, and culture-based methods, and culture independent (qPCR, metagenomics, and metatranscriptomics) were used to determine selective effects. The results from these experiments were varied.

Mostly, diclofenac, metformin, and 17-β-estradiol did not appear to strongly select for antibiotic resistance, metal and biocide resistance, nor significantly alter the community composition.

### One of three NADs selected for *intI1*

Only 17-β-estradiol selected for one of the three qPCR targets tested here (*intI1*), from 7µg/L to 5400µg/L, in a non-dose-dependent manner. This could suggest that the gene associated with the class I integron that is under selection provides a low level benefit to the community, but there is an upper limit to of this benefit. However, there is no indication of which genes associated with *intI1* may be driving this response. *IntI1* is often recommended as a gene to use in environmental AMR surveillance as it can be associated with a large range of ARGs (87), biocide, and metal resistance genes (60). Future research could sequence the gene cassettes within the class I integrons in these communities, to find those that are closest to the promoter (and therefore would be the most highly expressed). Regardless, selection for *intI1* suggests that there could be a risk of increased mobilisation of genes, which could be ARGs, and therefore increased potential for dissemination of AMR throughout 17-β-estradiol impacted bacterial communities. However, whether 17-β-estradiol can indeed increase HGT would need to be experimentally tested in future work.

However, interestingly, there was no selection for *cintI1* or *sul1* seen by 17-β-estradiol. This suggests that the gene/s under selection by 17-β-estradiol are not present on the clinical class I integrons. The more general (or environmental) class I integrons can sometimes contain no known ARGs, but can contain metabolic genes, that may be used to adapt to stress (88, 89). Therefore, potentially, the gene driving the response seen in *intI1* with 17-β-estradiol is not a gene typically involved in AMR.

### No broad scale effects on ARG relative abundances

Selection for class 1 integrons carrying metabolic/stress genes by 17-β-estradiol is further supported by the fact we did not observe any selection for ARGs by diclofenac, metformin, or 17-β-estradiol using metagenome analyses, and no increases in phenotypic resistance for the antibiotics tested. The presumptive coliforms in communities treated with 17-β-estradiol in fact increased in sensitivity to gentamicin and ampicillin. However, metformin appeared to select against *TolC,* part of the acrAB-TolC efflux pump, known to provide resistance to multiple antibiotics and disinfectants (56). Therefore, selection against *TolC* may reduce co-selection for AMR, however, this is not captured in the phenotypic analyses presented here. Metformin treatments of 13.6ug/L and greater appeared to decrease diversity of ARGs compared to the control, suggesting that metformin did select against specific ARGs, but that this could not be detected with statistical approaches used at the ARG or ARG class level.

### There were some changes to metal resistance gene abundances as indicated by metagenomic analysis

Diclofenac, metformin, and 17-β-estradiol did appear to alter relative abundances of some metal resistance genes, though often in a non-dose-dependent manner. After treatment with diclofenac, metformin, or 17-β-estradiol, the relative abundances of arsenic resistance genes (e.g. *arsA*), and mercury resistance genes (e.g. *merT*) were significantly altered (increased or decreased) across at least one concentration. This suggests that there could be a generalisable response, co-opting metal resistance pathways to manage any toxic effects of these NADs.

Treatment with 17-β-estradiol dose-dependently increased the relative abundance of three arsenic resistance genes (two versions of *arsB,* and *arsR*) and decreased the relative abundance of *ncrA*. The arsenic resistance genes are involved in the production of an arsenite efflux pump (90, 91). These genes were isolated (and annotated in BacMet) from three different bacterial (Gram positive and Gram negative) species - *Yersinia enterocolitica, Staphylococcus aureus*, and *Acidiphilium multivorum* (69). This suggests that the increase in these genes occurs across phyla, and the effect is not driven by a dominant species. This is supported by the lack of a significant alteration in the community composition following treatment with 17-β-estradiol. Potentially, this efflux pump can confer cross-resistance to 17-β-estradiol, and should be a target for future studies. The gene *ncrA* is involved in the ncrABCY efflux system (92) and its negative selection in this study suggests it is not beneficial to the community, potentially to offset the cost of positive selection for the arsenite efflux pump genes.

### Small unidentifiable changes to gene expression indicated by transcriptome analysis

There were no changes in ARGs that were expressed in response to diclofenac, metformin, or 17-β-estradiol, but there was decreased expression of various metal resistance genes, which is somewhat contradictory to increased relative abundances of these genes. For example, 17-β-estradiol reduced *arsR* expression at the highest concentration, but led to a dose-dependent increase in *arsR* abundance. Furthermore, this indicates that expression of genes is not always coupled to heritable selection of genes. Additionally, as mentioned, there were various gene transcripts that were only expressed in response to one of the three NADs that were not present in the control populations. These are uncharacterised, and it would be prudent for future work to identify these genes, in order to understand if these are related to antimicrobial resistance in some way, or are unknown genes involved in adaptation to pharmaceutical pollution, or general stress responses. However, since *arsR* appeared to be decreased in expression, but increased in absolute gene abundance, alongside an increase in *intI1* prevalence, this suggests that these genes are co-located with other unidentified resistance mechanisms. These mechanisms may be 17-β-estradiol specific, but further work would be required to identify these.

### Limitations

Of course, there are limitations associated with this work. Firstly, we used a mixed bacterial community derived from sewage and yet used experimental conditions that are unrealistic for this community (high temperature and high nutrient media). Therefore, in the environment or human gut, these effects may differ, since different species or strains may not be able to survive in our laboratory conditions that would survive in those environments. In order to address this, future work should aim to use environment-specific conditions (e.g. anaerobic conditions to mimic the human gut). This would aim to provide a more realistic set of experimental conditions, and allow us to understand whether anaerobic prokaryote responses would differ to the results presented here. Secondly, genes may have been selected for that confer low-levels of resistance to antibiotics or other antimicrobials but may not be identified as such in the ARGs-OAP database used in this work. Metabolic genes can confer antibiotic and biocide resistance (93) but would not be identified as ARGs. This is a common issue regarding environmental AMR work since databases used to identify ARGs are predominantly based on clinically derived data, and are therefore biased towards clinical pathogens and human associated bacteria. Thirdly, due to the nature of the experimental design, it was difficult to identify low effect size changes in the metagenome data. Community experiments are understandably noisy due to the large variation in strains and species present. Therefore, subtle changes in responses are difficult to identify. Further work should aim to include increased replicative power to try to be able to identify effects that may have been lost in this study.

Additionally, there were many unidentified genes that were identified to be associated with NAD only treatments in the transcriptome analysis. However, the current pipeline used was not able to identify these. Therefore, further work could aim to identify these, since they might indicate NAD specific effects on bacteria, and would help to understand the mechanism behind the inhibition of growth seen in this study.

Finally, we propose that these experimental conditions are not fully replicative of how bacterial communities would encounter environmental pollutants. In the environment, pollutants are present as a complex mixture, including NADs (23), antibiotics (94), and other pollutants, including chemicals used in agriculture (e.g. pesticides and insecticides) (95), biocides (e.g. those used in disinfection in hospital settings) (96), alongside chemicals from road run-off (97). Therefore, the effects of mixtures of these compounds might be more important in understanding the real-world effects of NADs. Even when prescribed clinically, NADs are often combined with other medication, whether transiently (e.g. with ad-hoc usage of painkillers) or long-term (e.g. multiple pharmaceuticals prescribed for one or multiple long-term conditions). Therefore, although the NADs tested here do not appear to co-select for antibiotic resistance when tested alone, what occurs when these NADs are in combination with one or multiple other compounds should be tested.

### Conclusion

This study suggests that diclofenac and metformin do not select for AMR in raw sewage derived bacterial communities. These data are similar to those presented recently by Hall et al., (28) which show that in *E. coli*, several NADs are capable of reducing growth, but do not co-select for antibiotic resistance. Here, we show this is also likely the case in a community model, which comprised of predominately *E. coli* bacteria. However, 17-β-estradiol did select for *intI1*, indicating this NAD could increase the mobilisation potential of ARGs. Further, our data indicates that metal resistance genes may be used by bacteria to counter stress caused by NAD pollution, and may be a co-selection pathway for both metal and antibiotic resistance, by selecting for components of multidrug efflux pumps. Future work should aim to understand these effects in both a human gut community (anaerobically), and in a freshwater experimental set up (including low temperature and limited-nutrient media), and to understand the effects on HGT after exposure to these NADs. This would indicate if the effects seen here in a wastewater community also occur at these different microbiomes.

## Authors’ contributions

AH designed the study, performed the experiments, performed bioinformatics analysis, and statistical analysis, and wrote and edited the manuscript. LZ, EF, JS, BKH provided comments for the final manuscript and writing. WHG designed the study, provided comments on interpretation and edited the manuscript. AKM designed the study, provided comments on interpretation, edited the manuscript, and was responsible for project administration. LZ, EF, JS, BKH, WHG, and AKM supervised the project.

All authors read and approved the final version.

## Competing interests

AKM and WHG have received funding for Collaborative Awards in Science and Engineering (CASE) PhD Studentships from the UK Government (UKRI) with CASE support from AstraZeneca. AKM has also previously advised the AMR Industry Alliance. The funders had no role in the conception nor writing of this paper.

## Acknowledgements

We thank the Exeter Sequencing Service for their work in doing the Illumina metagenomic sequencing. This project utilised equipment funded by the Wellcome Trust Institutional Strategic Support Fund (WT097835MF), Wellcome Trust Multi User Equipment Award (WT101650MA) and BBSRC LOLA award (BB/K003240/1).

## Data Availability Statement

All data associated with this paper will be available upon publication.

