## Supplementary Tables for "Antimicrobial effects, and selection for AMR by non-antibiotic drugs on bacterial communities"

**Supplementary Table 1. Plasmid associated genes that were significantly different in at least one treatment after exposure to diclofenac.**

| **Gene** | **Presumptive species** | **P value (*fdr* adjusted)** |
|---|---|---|
| *arsA* | *E. coli* | 0.00017 |
| *arsA* | *Acidiphilium multivorum* | <0.0001 |
| *arsA* | *E. coli* | 0.0011 |
| *arsB* | *E. coli* | <0.0001 |
| *arsB* | *S. aureus* | <0.0001 |
| *arsB* | *Staphylococcus xylosus* | <0.0001 |
| *arsB* | *E. coli* | <0.0001 |
| *arsB* | *Rhizobium meliloti* | <0.0001 |
| *arsB* | *A. multivorum* | <0.0001 |
| *arsC* | *S. aureus* | <0.0001 |
| *arsC* | *S. xylosus* | <0.0001 |
| *arsC* | *E. coli* | <0.0001 |
| *arsC* | *E. coli* | <0.0001 |
| *arsC* | *A. multivorum* | <0.0001 |
| *arsC* | *Rhizobium meliloti* | <0.0001 |
| *arsD* | *E. coli* | 0.012 |
| *arsD* | *E. coli* | 0.0039 |
| *arsR* | *Rhizobium meliloti* | <0.0001 |
| *arsR* | *E. coli* | <0.0001 |

| | | |
|---|---|---|
| *arsR* | *S. aureus* | <0.0001 |
| *arsR* | *E. coli* | <0.0001 |
| *arsR* | *S. xylosus* | <0.0001 |
| *arsR* | *E. coli* | <0.0001 |
| *merA* | *Bacillus cereus* | <0.0001 |
| *merA* | *Serratia marcescens* | 0.00066 |
| *merA* | *Pseudomonas sp. K-62* | 0.029 |
| *merA* | *Pseudomonas stutzeri* | <0.0001 |
| *merA* | *Thiobacillus ferrooxidans* | <0.0001 |
| *merB* | *Pseudomonas sp. K-62* | 0.00013 |
| *merB* | *S. marcescens* | 0.0026 |
| *merD* | *P. aeruginosa* | <0.0001 |
| *merD* | *P. stutzeri* | <0.0001 |
| *merD* | *P. stutzeri* | <0.0001 |
| *merD* | *S. marcescens* | <0.0001 |
| *merD* | *Ralstonia metallidurans* | <0.0001 |
| *merD* | *Pseudomonas sp. K-62* | <0.0001 |
| *merE* | Plasmid pDU1358 | 0.00083 |
| *merE* | *P. stutzeri* | 0.00024 |
| *merE* | *P. aeruginosa* | <0.0001 |
| *merP* | *S. marcescens* | 0.00014 |
| *merP* | *Pseudomonas sp. K-62* | <0.0001 |
| *merP* | *P. stutzeri* | <0.0001 |
| *merP* | *P. stutzeri* | <0.0001 |

| | | |
|---|---|---|
| *merR1* | *Pseudomonas sp. K-62* | <0.0001 |
| *merR2* | *Pseudomonas sp. K-62* | <0.0001 |
| *merR* | *S. marcescens* | <0.0001 |
| *merR* | *P. stutzeri* | 0.00088 |
| *merR2* | *P. stutzeri* | <0.0001 |
| *merT* | *Alcaligenes sp.* | <0.0001 |
| *merT* | *T. ferrooxidans* | <0.0001 |
| *merT* | *P. stutzeri* | <0.0001 |
| *merT* | *P. stutzeri* | <0.0001 |
| *merT* | *Pseudomonas sp. K-62* | <0.0001 |
| *merT* | *S. marcescens* | <0.0001 |
| *ncrA* | *Hafnia alvei* | 0.015 |
| *ncrC* | *Enterobacter cloacae subsp. cloacae* | 0.022 |
| *qacEdelta1* | *P. aeruginosa* | 0.015 |
| *terC* | *Alcaligenes sp.* | 0.029 |
| *terD* | *Alcaligenes sp.* | 0.042 |

**Supplementary Table 2. Plasmid associated genes that were significantly different in at least one treatment after exposure to metformin**

| Gene | Presumptive species | P value (*fdr* adjusted) |
|---|---|---|
| *arsA* | *E. coli* | <0.0001 |
| *arsA* | *A. multivorum* | <0.0001 |
| *arsA* | *E. coli* | <0.0001 |
| *arsB* | *E. coli* | <0.0001 |
| *arsB* | *S. aureus* | <0.0001 |
| *arsB* | *St. xylosus* | <0.0001 |
| *arsB* | *E. coli* | <0.0001 |
| *arsB* | *R. meliloti* | <0.0001 |
| *arsB* | *A. multivorum* | <0.0001 |
| *arsC* | *S. aureus* | <0.0001 |
| *arsC* | *E. coli* | <0.0001 |
| *arsC* | *E. coli* | <0.0001 |
| *arsC* | *A. multivorum* | <0.0001 |
| *arsC* | *R. meliloti* | <0.0001 |
| *arsR* | *R. meliloti* | 0.00017 |
| *arsR* | *E. coli* | <0.0001 |
| *arsR* | *E. coli* | 0.011 |
| *arsR* | *E. coli* | <0.0001 |
| *merA* | *B. cereus* | <0.0001 |

| | | |
|---|---|---|
| *merA* | *S. marcescens* | 0.0056 |
| *merA* | *Pseudomonas sp. K-62* | 0.022 |
| *merA* | *S. aureus* | 0.0065 |
| *merA* | *P. stutzeri* | 0.00064 |
| *merA* | *T. ferrooxidans* | 0.0014 |
| *merB* | *S. marcescens* | 0.0018 |
| *merD* | *P. aeruginosa* | <0.0001 |
| *merD* | *P. stutzeri* | <0.0001 |
| *merD* | *P. stutzeri* | <0.0001 |
| *merD* | *S. marcescens* | <0.0001 |
| *merD* | *R. metallidurans* | <0.0001 |
| *merD* | *Pseudomonas sp. K-62* | <0.0001 |
| *merE* | Plasmid pDU1358 | <0.0001 |
| *merP* | *S. marcescens* | 0.013 |
| *merP* | *Pseudomonas sp. K-62* | <0.0001 |
| *merP* | *P. stutzeri* | 0.017 |
| *merP* | *P. stutzeri* | 0.00015 |
| *merR1* | *Pseudomonas sp. K-62* | <0.0001 |
| *merR2* | *Pseudomonas sp. K-62* | <0.0001 |
| *merR* | *S. marcescens* | 0.0080 |
| *merR* | *P. stutzeri* | 0.026 |
| *merR2* | *P. stutzeri* | <0.0001 |
| *merR* | *T. ferrooxidans* | 0.0095 |
| *merT* | *Alcaligenes sp.* | 0.00013 |

| | | |
|---|---|---|
| *merT* | *T. ferrooxidans* | <0.0001 |
| *merT* | *P. stutzeri* | <0.0001 |
| *merT* | *P. stutzeri* | <0.0001 |
| *merT* | *Pseudomonas sp. K-62* | <0.0001 |
| *merT* | *S. marcescens* | <0.0001 |
| *ncrB* | *H. alvei* | 0.033 |

**Supplementary Table 3. Plasmid related metal and biocide resistance genes that are significantly different in at least one treatment in populations evolved with 17-β-estradiol**

| Gene | Presumptive species | P value (*fdr* adjusted) |
|---|---|---|
| *arsA* | *E. coli* | <0.0001 |
| *arsA* | *A. multivorum* | <0.0001 |
| *arsA* | *E. coli* | 0.00012 |
| *arsB* | *E. coli* | 0.012 |
| *arsB* | *S. aureus* | <0.0001 |
| *arsB* | *S. xylosus* | <0.0001 |
| *arsB* X | *E. coli* | <0.0001 |
| *arsB* | *R. meliloti* | <0.0001 |
| *arsB* | *A. multivorum* | <0.0001 |
| *arsC* | *S. aureus* | <0.0001 |
| *arsC* | *S. xylosus* | <0.0001 |
| *arsC* | *E. coli* | <0.0001 |
| *arsC* | *E. coli* | <0.0001 |
| *arsC* | *A. multivorum* | <0.0001 |
| *arsC* | *R. meliloti* | <0.0001 |
| *arsD* | *E. coli* | 0.0057 |
| *arsR* | *R. meliloti* | 0.0079 |
| *arsR* | *E. coli* | <0.0001 |

| | | |
|---|---|---|
| *merA* | *S. marcescens* | <0.0001 |
| *merA* | *P. stutzeri* | <0.0001 |
| *merA* | | 0.0055 |
| *merB* | *S. marcescens* | 0.00081 |
| *merD* | *P. aeruginosa* | <0.0001 |
| *merD* | *P. stutzeri* | <0.0001 |
| *merD* | *P. stutzeri* | <0.0001 |
| *merD* | *S. marcescens* | <0.0001 |
| *merD* | *R. metallidurans* | <0.0001 |
| *merD* | *Pseudomonas sp. K-62* | <0.0001 |
| *merE* | *Plasmid pDU1358* | <0.0001 |
| *merE* | *P. stutzeri* | 0.013 |
| *merE* | *P. aeruginosa* | <0.0001 |
| *merP* | *S. marcescens* | 0.00016 |
| *merP* | *Pseudomonas sp. K-62* | 0.0016 |
| *merP* | *P. stutzeri* | 0.0031 |
| *merP* | *P. stutzeri* | 0.010 |
| *merR1* | *Pseudomonas sp. K-62* | <0.0001 |
| *merR2* | *Pseudomonas sp. K-62* | <0.0001 |
| *merR* | *S. marcescens* | 0.012 |
| *merR* | *P. stutzeri* | 0.039 |
| *merR2* | *P. stutzeri* | <0.0001 |
| *merR* | *T. ferrooxidans* | 0.0062 |
| *merT* | *Alcaligenes sp.* | 0.00046 |

| | | |
|---|---|---|
| *merT* | *T. ferrooxidans* | <0.0001 |
| *merT* | *P. stutzeri* | 0.011 |
| *merT* | *P. stutzeri* | <0.0001 |
| *merT* | *Pseudomonas sp. K-62* | <0.0001 |
| *merT* | *S. marcescens* | 0.0093 |
| *ncrA* | *Leptospirillum ferriphilum* | 0.018 |
| *ncrA* | *H. alvei* | 0.0019 |
| *ncrB* | *L. ferriphilum* | 0.0045 |
| *ncrC* | *Enterobacter cloacae subsp. cloacae* | 0.029 |
| *ncrC* | *S. marcescens* | 0.024 |
