## Supplementary Figures for "Antimicrobial effects, and selection for AMR by non-antibiotic drugs on bacterial communities"

**Supplementary File 1**

**Figure 1. QPCR prevalence of A) *intI1*, B) *cintI1*, and C) *sul1* as a function of diclofenac concentration. 5 biological replicates per treatment.**

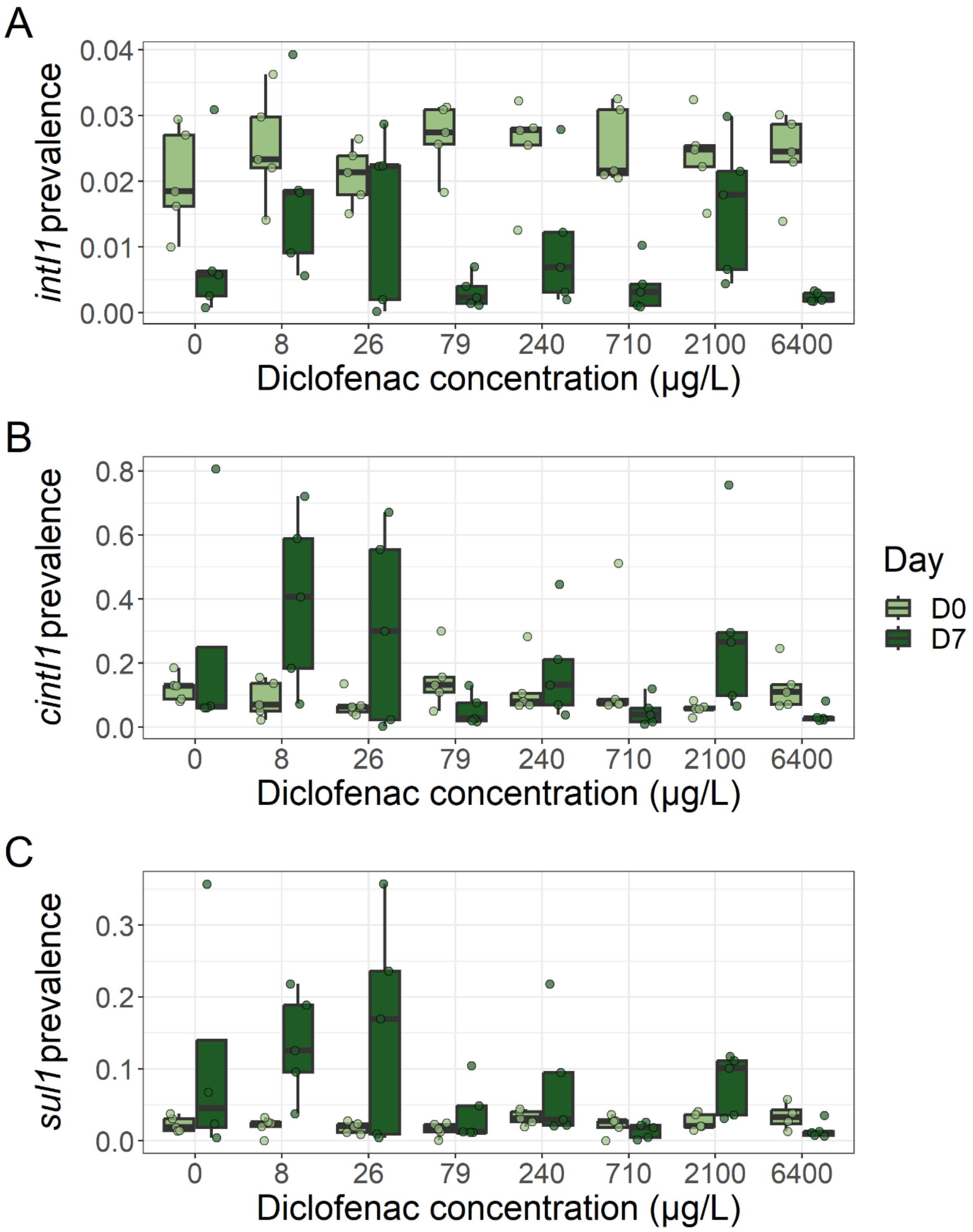

**Figure 2. QPCR prevalence of A) *intI1*, B) *cintI1*, and C) *sul1* as a function of metformin concentration. 5 biological replicates per treatment.**

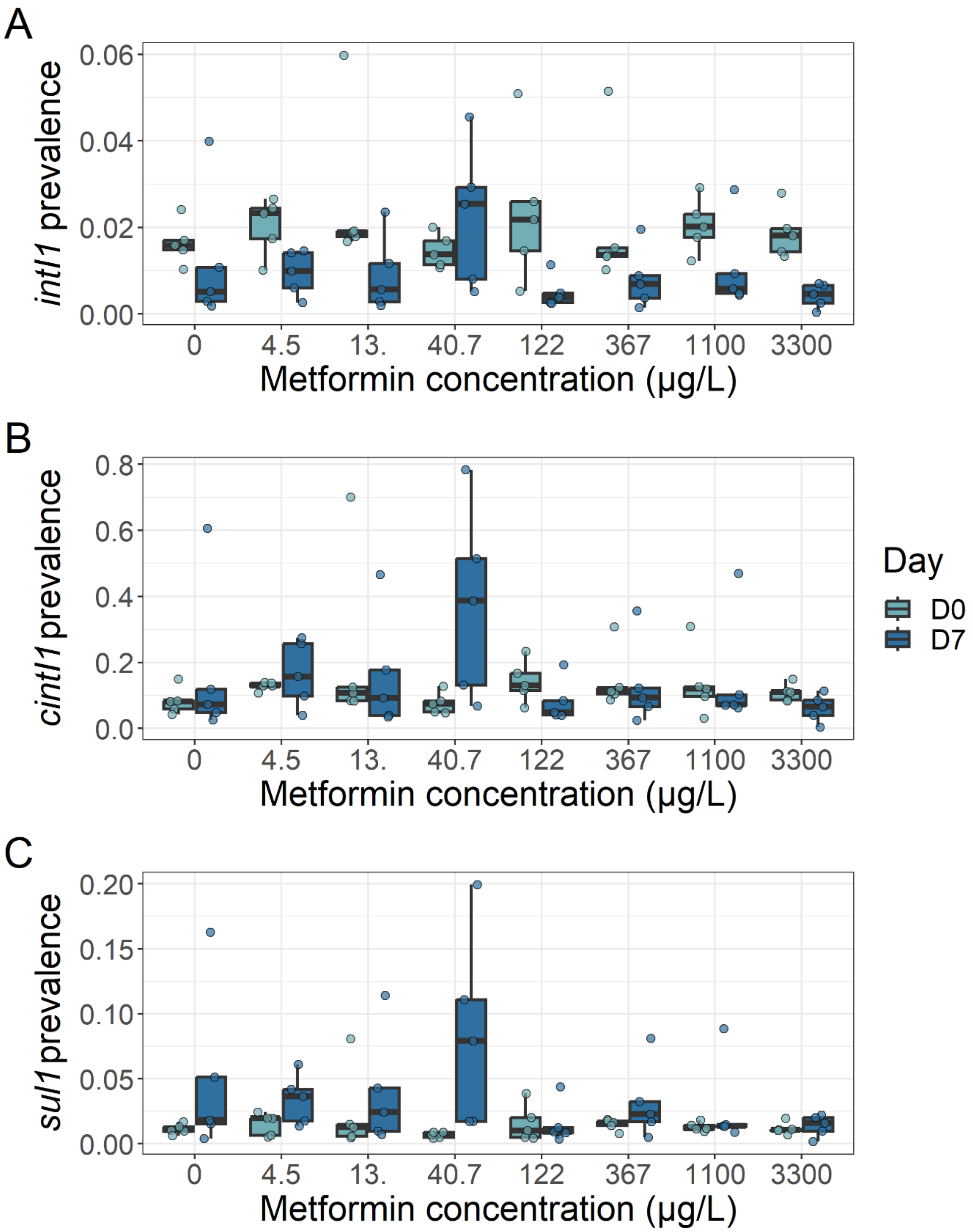

**Figure 3. QPCR prevalence of A) *cintI1*, and B) *sul1* as a function of 17-β-estradiol concentration. 5 biological replicates per treatment.**

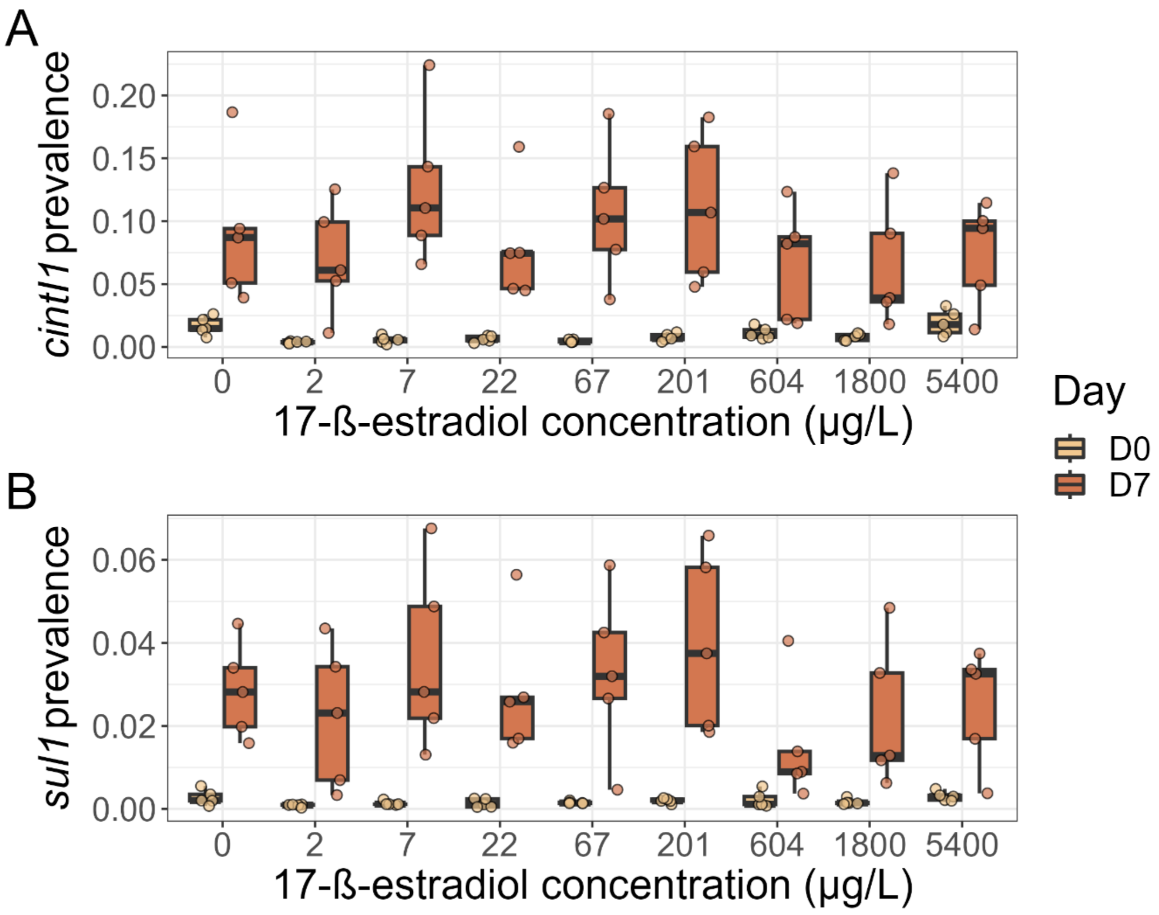

**Figure 4. Heat map of relative abundance of bacterial species after seven-day evolution with diclofenac.**

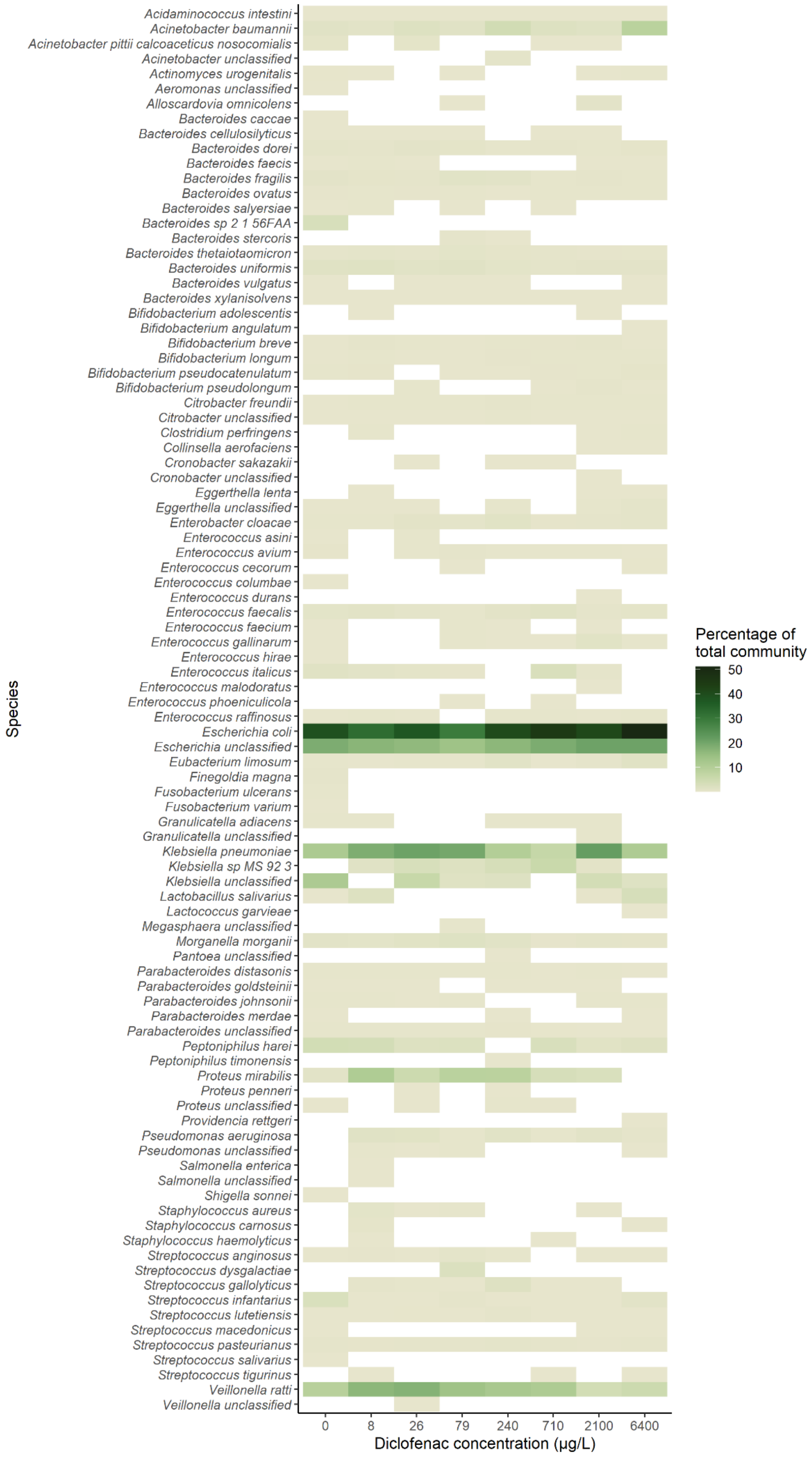

**Figure 5. Heat map of relative abundance of bacterial species after seven-day evolution with metformin. Evolved communities shown only.**

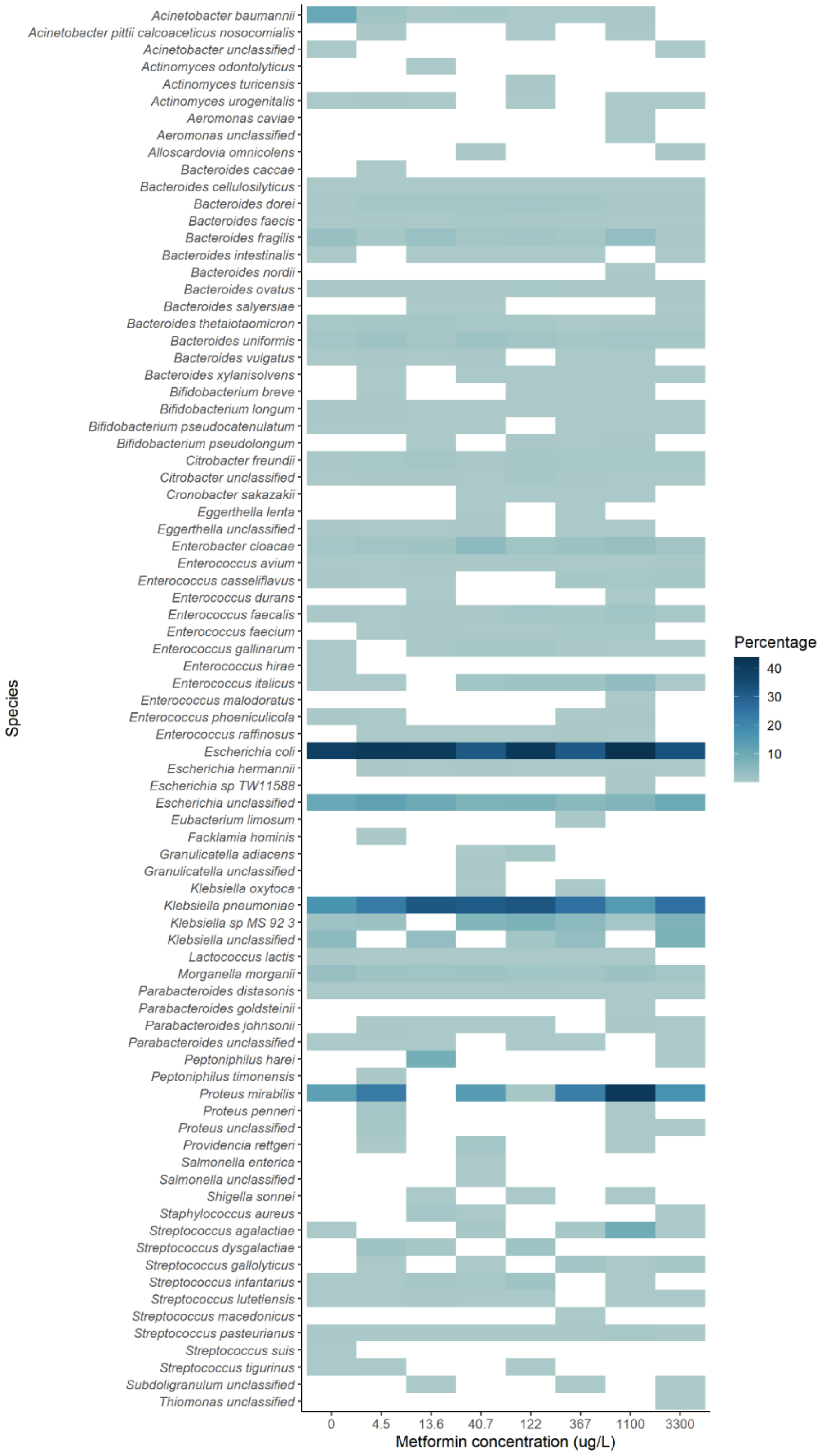

**Figure 6. Heat map of relative abundance of bacterial species after seven-day evolution with 17-β-estradiol. Evolved communities shown only**

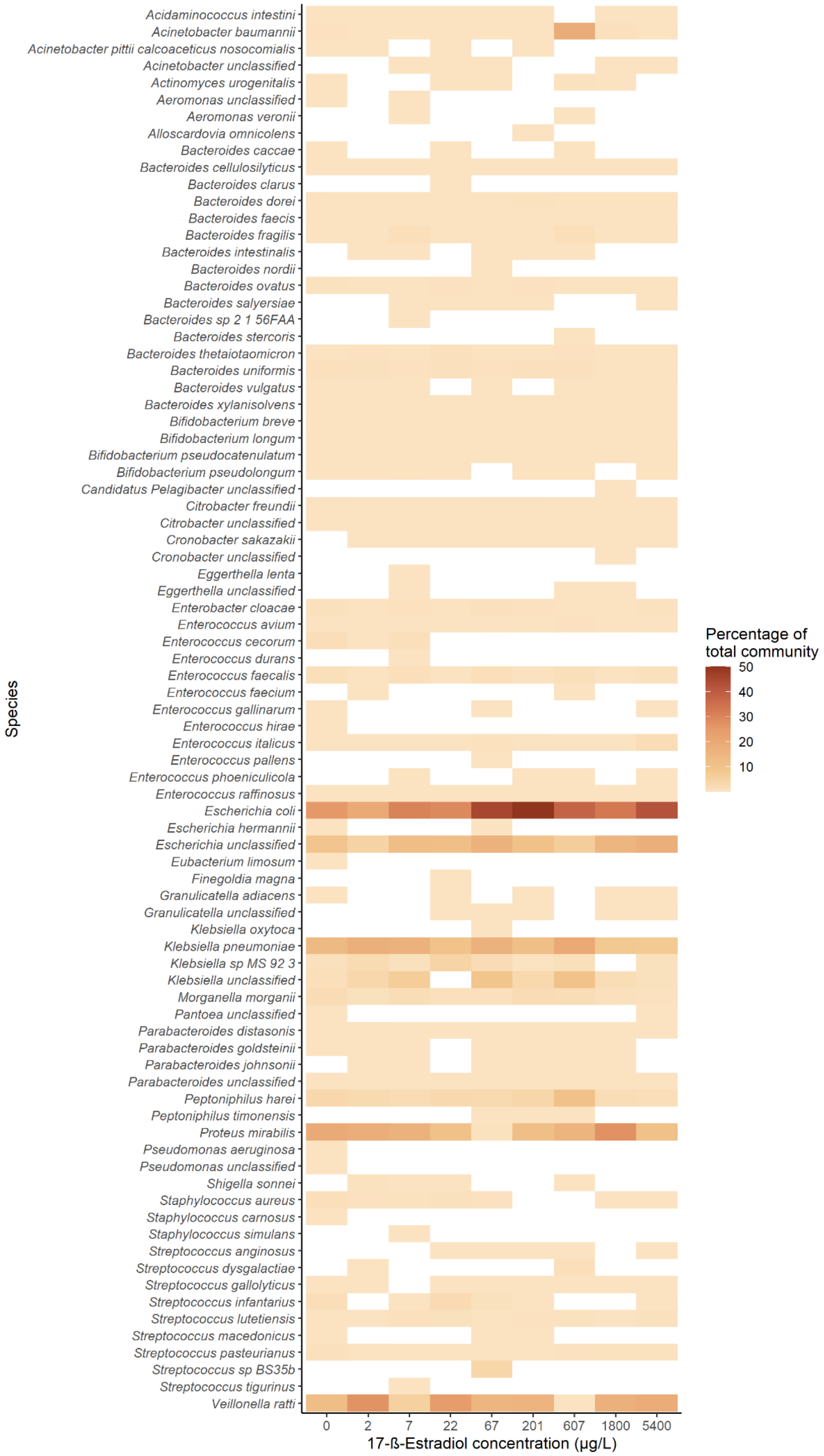

**Figure 7. Relative abundance of ARGs in communities evolved with diclofenac. Abundance of genes are normalised to 16S rRNA reads. Not showing multidrug resistance genes, and non-classified genes, since removal of these for the heatmap allowed for better visualisation of all classes due to the high abundance of multidrug and unclassified genes.**

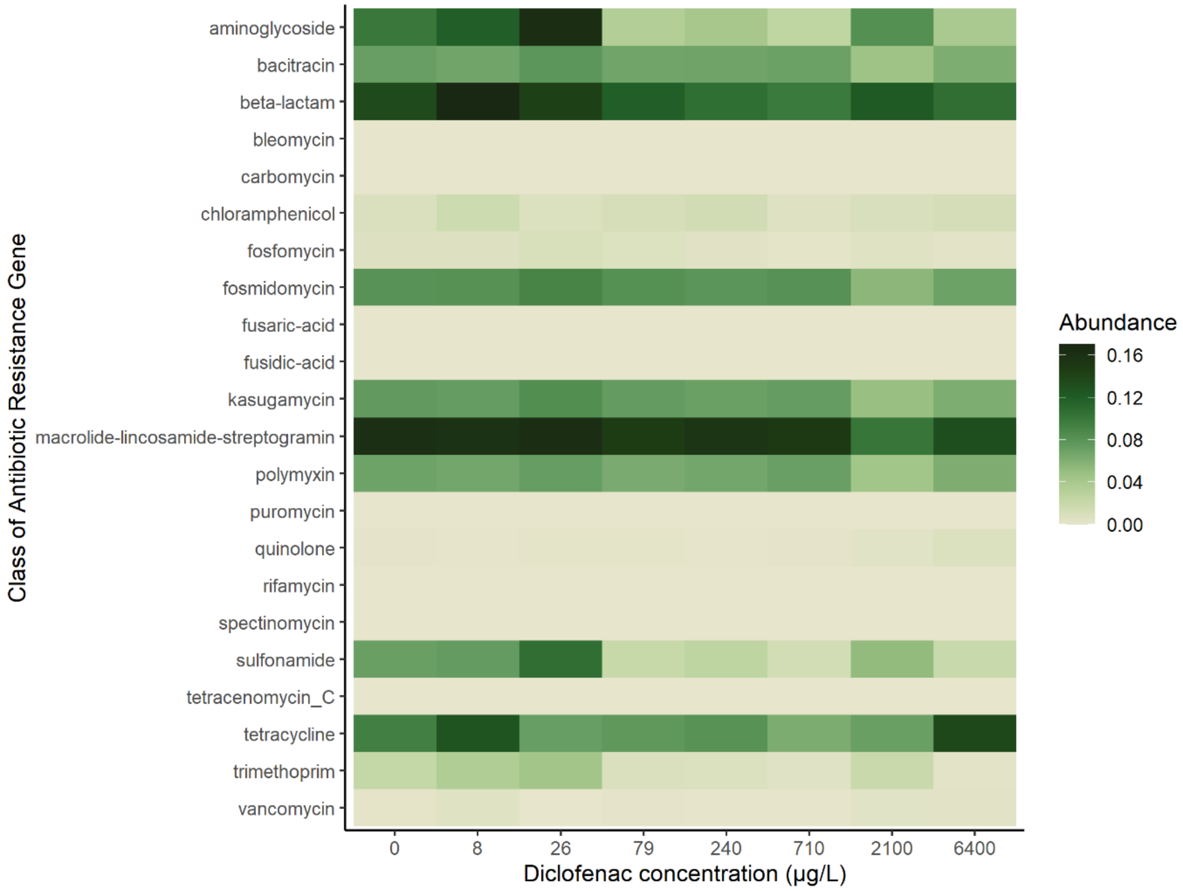

**Supplementary Figure 8. Abundance of genes are normalised to 16S rRNA reads. Not showing multidrug resistance genes, and non-classified genes since removal of these for the heatmap allowed for better visualisation of all classes due to the high abundance of multidrug and unclassified genes.**

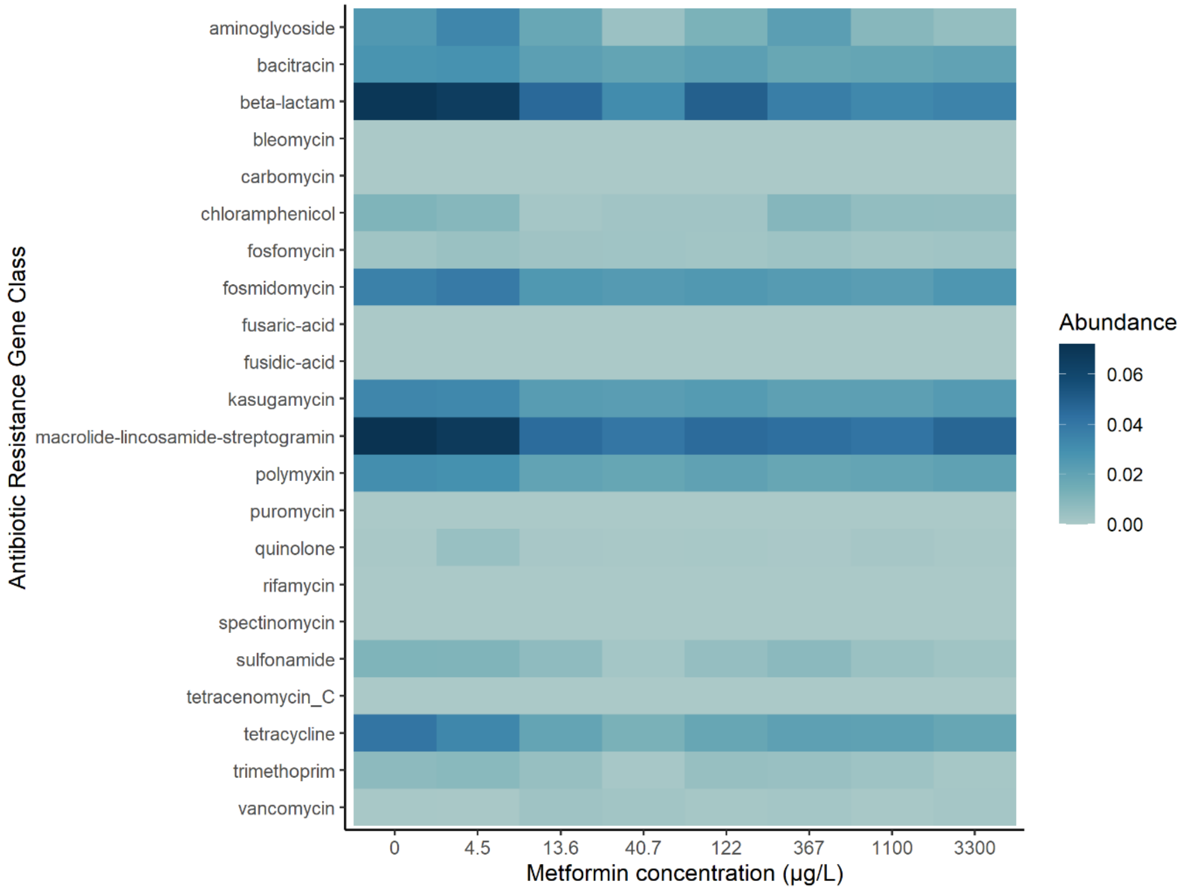

**Supplementary Figure 9. Relative abundance of ARGs per gene class after exposure to 17-β-estradiol. Multidrug resistance genes, and genes that are unclassified in the database are not shown since removal of these for the heatmap allowed for better visualisation of all classes due to the high abundance of multidrug and unclassified genes.**

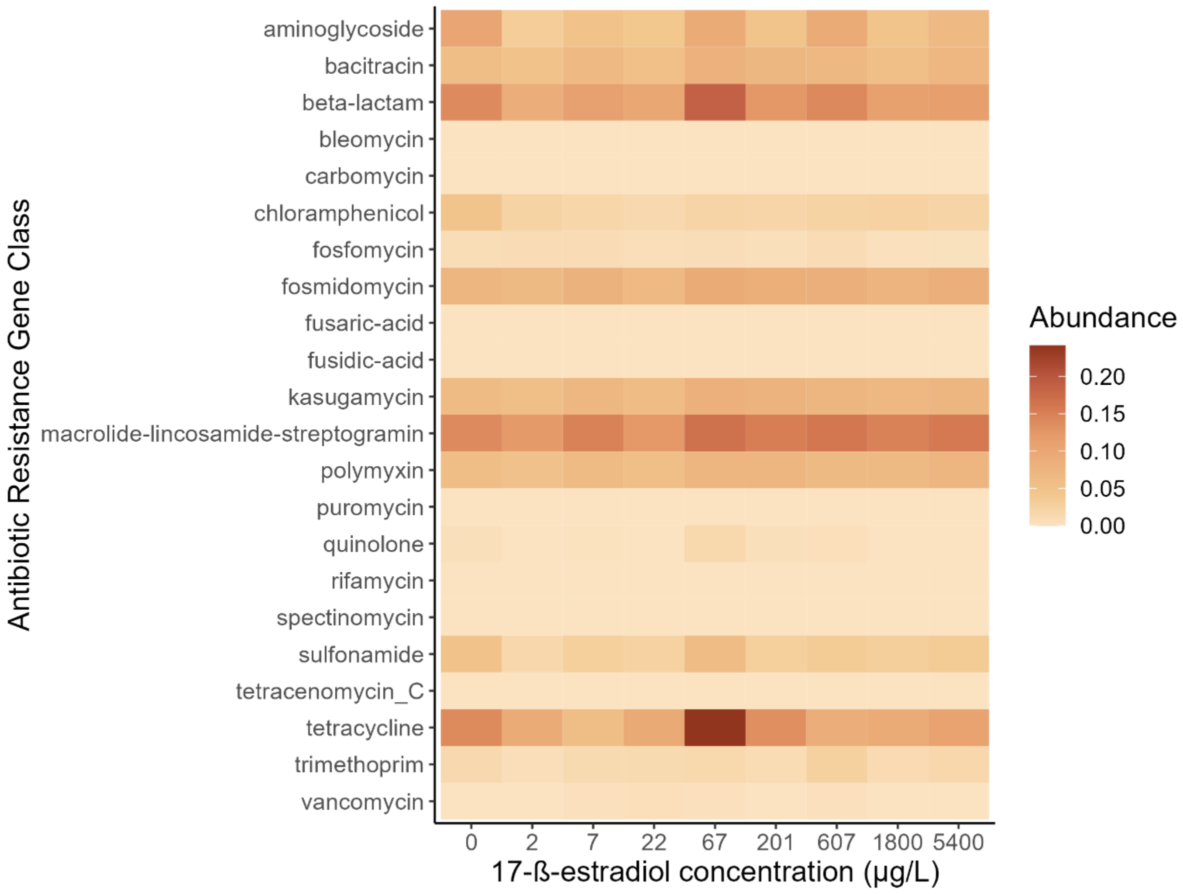
